# Rapid evolution of pesticide resistance via adaptation and interspecific introgression in a major North American crop pest

**DOI:** 10.1101/2023.10.04.560869

**Authors:** Henry L. North, Zhen Fu, Richard Metz, Matt A. Stull, Charles D. Johnson, Xanthe Shirley, Kate Crumley, Dominic Reisig, David L. Kerns, Todd Gilligan, Tom Walsh, Chris D. Jiggins, Gregory A. Sword

## Abstract

Insect crop pests threaten global food security. This threat is amplified through the spread of non-native species and the evolution of pesticide resistance, which can be introduced to a population though *de novo* mutation or gene flow. We investigate these processes in an economically important noctuid crop pest, *Helicoverpa zea*, which has rapidly evolved resistance to several pesticides. Its sister species *Helicoverpa armigera*, first detected as an invasive species in Brazil in 2013, introduced the pyrethroid resistance gene *CYP337B3* to South American *H. zea* via introgression. To understand whether this contributes to pesticide resistance in North America, we sequenced 237 *H. zea* genomes across 10 sample sites in the US. First, we report *H. armigera* introgression into the North American *H. zea* population. Two individuals sampled in Texas in 2019 carry *H. armigera* haplotypes in a 4Mbp region containing *CYP337B3*. Second, we show that the remarkable dispersal ability of *H. zea* results in a panmictic North American population. Third, we detect signatures of selection in non-admixed *H. zea*, identifying a selective sweep at a second pesticide resistance locus with a similar name: *CYP333B3*. We estimate that its derived allele conferred a ∼4.9% fitness advantage and show that this estimate explains independently observed rare nonsynonymous *CYP333B3* mutations approaching fixation over a ∼20-year period. We also detect putative signatures of selection at a kinesin gene associated with Bt resistance. Our results show that pesticide resistance in *H. zea* evolved rapidly and recently via two independent mechanisms: interspecific introgression and rapid intraspecific adaptation.

## Introduction

Insect pests destroy 5-20% of major grain crop production, and losses are set to increase substantially over coming decades as a result of climate change, the evolution of pesticide resistance, and the spread of invasive species via international trade routes (Deutsch et al. 2018; Gould et al. 2018; Paini et al. 2016). Understanding the role that evolution plays in the ecology of pests and their outbreaks is therefore a priority (Luke et al. 2023; Karlsson Green et al. 2020). The short generation times and large effective population sizes of many invertebrate pest species results in high rates of molecular evolution and rapid allele frequency shifts in response to natural selection (Thomas et al. 2010; Petit & Barbadilla 2009). At the same time, extreme selective regimes imposed by pesticide exposure can result in rapid adaptation (Hawkins et al. 2019). The strong dispersal ability of many insect pests, especially in the context of contiguous habitat in monoculture, means that pest populations often exist as highly connected metapopulations that cover agricultural landscapes (Mazzi & Dorn 2012). As a consequence, fitness-enhancing alleles can readily spread across space (McDonald & Linde 2003). Human activity can also mediate dispersal at larger geographic scales (*i.e.*, across agricultural systems and continents) such that insecticide resistance can rapidly arise through gene flow – a process significantly faster than adaptation from de-novo mutation (Tay & Gordon 2019). The extent of global trade networks means that many closely related pest species come into secondary contact, opening up the possibility of adaptive introgression not only between populations, but also between divergent ecotypes and species (Valencia-Montoya et al. 2020; Song et al. 2011). Together these factors enhance the evolutionary potential of insect pests, with two key consequences. First, adaptive responses can occur at timescales relevant to year-to-year pest management strategies. Second, the global connectedness of many pest populations means that such strategies must be multilateral. Large-scale genomic monitoring is widely accepted as a promising emerging means of informing management action to address these consequences.

Population genetics has long been used as a tool for quantifying evolutionary change in agricultural pests, especially with respect to insecticide resistance (Mallet 1989). The additional information and contiguity of resolution offered by genome-resequencing data have created renewed interest in this field, and genomic approaches are clearly emerging as a key tool for monitoring pest populations under both proactive and reactive management plans (Sherpa & Després 2021; Hamelin & Roe 2020; Neafsey et al. 2021; North et al. 2021). Recent studies have demonstrated the use of population genomics approaches to define management units by quantifying population connectivity (Chen et al. 2021; Paris et al. 2022), identifying loci associated with the evolution of pesticide resistance by inferring the action of selection (Love et al. 2023), and reconstructing the spatial spread of species or alleles of interest (Tay et al. 2022). With appropriate analysis and sampling design, population genomics can be used to extract otherwise-inaccessible biological information to understand the evolutionary history of pest populations and inform management plans.

*Helicoverpa zea*, commonly known as the corn earworm, is a polyphagous noctuid moth common throughout the Americas. A notorious pest of maize and cotton, *H. zea* is one of the most economically significant crop pests in the agricultural powerhouses of Brazil and the United States (Olivi et al. 2019; Fitt 1989; Cook & Threet 2019; Olmstead et al.; Cunningham & Zalucki 2014). Although maize is its dominant host plant, larvae are known to feed on at least 122 other species, of which 29 are major crops including wheat, soy, rice, sorghum, and tomato (Cunningham & Zalucki 2014). Larvae tend to feed on the fruiting body of the plant, thereby directly damaging produce (Luttrell & Jackson). Generation times vary depending on latitude (5-10 generations/yr at lower altitudes), though facultative diapause enables pupa to persist underground up to at least 40°N during winter (Hardwick 1965; Morey 2010; Parajulee et al. 2004). Adults are highly effective long-distance dispersers, expanding northward into extensive areas of maize production during summer to a latitude of ∼52°N in flights large enough to detect using ground-based radar (Jones et al. 2018; Westbrook 2008). The species’ range is expected to expand twofold by 2099 as warmer winters reduce the number of lethal low-temperature events (Lawton et al. 2022). Repeated admixture due to seasonal re-establishment from southern populations, combined with long-range dispersal over highly connected agricultural habitat, result in a highly connected and genetically diverse metapopulation in the north (Seymour et al. 2016; Margosian et al. 2009).

Multiple studies have found that the local distribution of crops producing *Bacillus thuringiensis* (Bt) toxins in a given year can predict *H. zea* damage in subsequent years (Arends et al. 2021, 2022). This observation implies that selection results in geographically localised phenotypic change between generations, so parent-offspring dispersal should primarily occur at the same geographic scale. In contrast, population genetics studies have reported effective panmixia across the North American range, though there is mixed evidence for this observation, and the only study to employ whole-genome data compared just two sample sites (Perera et al. 2020; Seymour et al. 2016; Taylor et al. 2021). Increased sampling effort — in terms of both geographic range and number of loci — can reveal population structure that is otherwise undetectable, as demonstrated in studies of *Helicoverpa armigera* (Jin et al. 2023; Zhang et al. 2022). Characterising the landscape of effective migration is key to understanding how rapidly adaptive variants underlying pesticide resistance can spread.

*H. zea* has evolved resistance to several pesticides. In the United States, the organochloride DDT was effective for *H. zea* control from its post-war implementation until the 1960s; resistance to methyl parathion (an organophosphate) introduced 1960s was initially detected in the 1970s; several pyrethroids introduced in the 1970s were effective until resistance started to become apparent in the 1990s and 2000s (Abd-Elghafar et al. 1993; Walsh et al. 2022). The specific mode of action of these pesticides means that resistance phenotypes could in many cases be underlain by mutations at one or few loci and therefore evolve rapidly in response to strong selection (Ibrahim et al. 2015). As a member of the ‘megapest’ genus of polyphagous herbivores, *Helicoverpa*, *H. zea* may be particularly well-equipped to evolve resistance due to its ecology and its capacity to metabolise a wide array of host plant defences (Gordon et al. 2010; Mallet 1989; Bras et al. 2022). Although many phenotypes with different genetic architectures can underly resistance to a given pesticide, certain gene families have been repeatedly implicated as targets of selection. This is true of cytochrome P450 genes involved in xenobiotic metabolism not only in *Helicoverpa* but in pest species spanning the tree of life (Kreiner et al. 2019; Nauen et al. 2022).

By the 1990s it became clear that the evolution of resistance was outpacing the development of novel pesticides, highlighting the need for a strategic shift toward integrated pest management. New approaches implemented for lepidopteran pests included the use of cotton and maize crops engineered to produce Bt toxins, which today constitute 82% of US maize crops and 88% of cotton (Reisig et al. 2022). In contrast to the nerve- and muscle-targeting insecticides discussed above, Bt toxins induce pore formation in the midgut membrane. Resistance to these toxins therefore requires selection on an entirely different set of loci. The most successful management plans made use of Bt crops expressing multiple toxins (*i.e.*, different Cry or Vip proteins) at high concentrations (>25x the dose required to kill susceptible insects) planted among non-Bt refuges in which rare resistant individuals could reproduce with susceptible mates. For insect pests generally, this approach was broadly successful at minimising unidirectional selection pressures, reducing net pesticide use and slowing the rate of resistance evolution. However, *H. zea* is among a few pest species to have evolved Bt resistance in the field. Cry1Ac resistance was reported in the early 2000s, at which point multi-toxin crops additionally expressing Cry2 were planted (Ali et al. 2006). By 2016, resistance to Cry1 toxins had become common throughout the US and Cry2 resistance was emerging (Dively et al. 2016; Reisig et al. 2018). Recent studies concluded that resistance to Cry2Ab2 has become common, and that selection was ongoing as of 2019 (Yu et al. 2021; Huang et al. 2023). The genetic basis of Bt resistance is known to be complex and likely arose from standing variation, with unique genetic architectures underlying resistance to different Cry toxins (Taylor et al. 2021; Benowitz et al. 2022).

In addition to intraspecific adaptation, a major concern for the spread of pesticide resistance in North American *H. zea* is through interspecific introgression from its sister species *H. armigera*. Commonly known as the cotton bollworm, *H. armigera* has a broad Afro-Eurasian native range. Among the most economically damaging crop pests in the world, *H. armigera* is more polyphagous and resistant to a substantially broader array of pesticides compared with *H. zea* (Cunningham & Zalucki 2014). *H*. *armigera* was first detected in Brazil in 2013, where the two species hybridized (Tay et al. 2013; Anderson et al. 2018; Ivey & Hillier 2023). This resulted in the adaptive introgression of the *CYP337B3* gene into South American *H. zea* populations (Valencia-Montoya et al. 2020). *CYP337B3*, otherwise absent in *H. zea*, is the result of unequal crossover from two other cytochrome P450 genes and confers fenvalerate resistance (Joußen et al. 2012). The variant has arisen multiple times in *H. armigera*, and can encode resistance to various pyrethroids including cypermethrin and deltamethrin (Durigan et al. 2017; Rasool et al. 2014)

*H. armigera* is now established throughout much of South and Central America. The risk of *H. armigera*, or admixed *H. armigera-zea* individuals, spreading into suitable North American habitat is considered high, and such an event would put at risk US crop production valued at $USD 78 billion per annum (Kriticos et al. 2015). *H. armigera* has been intercepted at US ports more than 1000 times, suggesting that introduction via shipping routes from its broad Afro-Eurasian range is also a substantial risk (Kriticos et al. 2015). Since the two species are difficult to distinguish phenotypically, and because of the extent of interspecific admixture in South America, the detection of invasive *H. armigera* alleles into North requires genetic surveillance of native *H. zea* populations. To date, there have been no published reports in the scientific literature of *H. armigera* establishing on the North American mainland. In 2015, three specimens captured in Florida carried the *H. armigera* COI haplotype (Tembrock et al. 2019), and multiple adults were captured in areas adjacent to Chicago O’Hare International Airport (USDA APHIS), though these are isolated incidents. There have been unpublished reports of *CYP337B3* detected in *H. zea* survey samples, though this observation may result from parallel evolution, as observed in *H. armigera* (Rasool et al. 2014).

Here, we use a population genomics approach to (*i*) test for signatures of introgression from *H. armigera* into North American *H. zea*, (*ii*) characterise effective migration across space to understand how rapidly resistance alleles may spread, and (*iii*) conduct a genome-wide scan for evidence of selective sweeps at known pesticide and Bt resistance loci. To achieve this, we resequenced the genomes of 237 *H. zea* individuals across 10 locations in the US collected in 2019 (see Methods).

## Results

### Evidence for introgression of pesticide resistance genes into North American H. zea

We tested for the presence of *H. armigera* ancestry by calculating 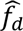 20kbp windows for individuals from each sample, using *H. zea* samples collected in Louisiana in 2002 as representative non-admixed samples and *H. punctigera* as an outgroup. We repeated this test for hybrids sampled in Brazil used as a positive control. The distribution of 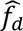 was centred on zero for all test sets apart from the positive control (Figure 2A). The same result is seen in samples reported in Taylor *et al*. (2021), collected in the US in 2012 and 2017. These results indicate balanced proportions of ABBA and BABA patterns, indicating little to no introgression at any sample site, at least compared to the magnitude of introgression seen in Brazil. However, 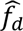 is elevated on chromosome 15 among samples collected in 2019 in Jackson County, TX (Figure 2B). Computing 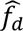 for each of the 16 samples from this site individually shows that the pattern is driven by extreme values of 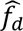 in only two individuals (Supplementary Figure S1).

For those individuals, 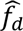 peaks at the *CYP337B3* locus (Figure 3). Nucleotide variation for these individuals is reduced across most of Chromosome 15 relative to *H. armigera*, but matches *H. armigera* in a terminal ∼4Mbp region around *CYP337B3*. Given that the effective population size of *H. armigera* is twice that of *H. zea* (Anderson et al. 2018), this pattern suggests *H. armigera* ancestry dominates in this region. Based on patterns 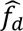 and *π*, we defined two segments of chromosome 15: A (0-9Mbp), which shows no signs of admixture, and B (>9Mbp) in which *H. armigera* haplotypes have introgressed.

We reasoned that if segment B consists of largely *H. armigera* ancestry, genetic differentiation and divergence from *H. armigera* should be lower in this region. Segment B shows reduced genetic differentiation relative to *H. armigera* (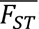 = 0.11 compared to 0.05 in Segment A; *p*<0.01, t = 25.606, df = 656, Welsch two-sample T test; Figure 3C). Genetic divergence also differed by segment (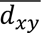 = 0.086 and 0.073 respectively; *p*<0.01, t = 10.495, df = 515.7; Supplementary Figure S2); this this difference was less pronounced, as expected given that differentiation between species should accumulate slower than differentiation.

These differences were apparent when visualising the data with principal components analysis (PCA). At segment A, the two admixed individuals cluster completely within other *H. zea* samples from 2019 in principal component space, whereas at segment B the individuals are closer to known Brazilian hybrid samples (Figure 4).

We next sought to determine whether the admixed samples carried the *H. armigera CYP337B3* gene, and if so, to use the gene tree to identify a potential *H. armigera* source population. This is because *CYP337B3* has arisen independently in multiple *H. armigera* populations. To investigate this, we reconstructed a maximum likelihood gene tree at the *CYP337B3* locus (HaChr13:11436565-11440168, as mapped by Anderson *et al*. (Anderson et al. 2018) and used by Valencia Montoya *et al*. (Valencia-Montoya et al. 2020)), comparing the admixed samples to publicly available *H. armigera* samples representing the breadth of the species’ phylogeographic diversity. The two admixed individuals form a clade with *H. armigera* samples at the *CYP337B3* locus (Figure 5). In 100% of bootstrap iterations, non-admixed *H. zea* samples were split from the clade consisting of admixed individuals and *H. armigera* samples. However, bootstrap support values were low within the *H. armigera* clade; there was no resolution to distinguish source populations. This result is consistent with low population structure within *H. armigera* and directional selection at this locus across the species’ native and invasive range (Ni et al. 2023; Jin et al. 2023).

Together, these results suggest that two of 237 individuals sampled are admixed, that introgression in these two samples is concentrated around the *CYP337B3* locus, and that both samples carry at least one *H. armigera CYP337B3* allele. Through its semidominant mode of action both individuals are therefore resistant to fenvalerate (Joußen et al. 2012).

### High connectivity across the North American *H. zea* metapopulation

We next sought to characterise the extent of allele sharing among non-admixed samples across the landscape to better understand how rapidly adaptive alleles (including those introduced from *H. armigera*) could spread. There was no positive relationship between geographic distance and genetic differentiation between sampling sites (one-sided Pearson’s product moment correlation for genome-wide average *F_ST_*: *t* = −2.8132, 43 degrees of freedom, *p* = >0.05), and the overall level of genetic differentiation was very low between sample sites (*F_ST_* < 0.01), so the North American *H. zea* metapopulation shows no signs of isolation by distance (Figure 6). In effect, individuals collected at all sample sites have the properties of a single Wright-Fisher-like population, meaning that segregating sites among samples from all locations could be used to detect selective sweeps. This provided us with substantial statistical power for our third aim.

### Signatures of selection throughout the genome

Given the absence of population structure among our samples, we included all individuals in a genome-wide scan for selective sweeps. We identified putative sweeps as extreme outliers for the composite likelihood ratio (CLR) implemented in *SweepFinder2* (see Methods). Three sweeps occurred on chromosomes 13, 15 and 25 (Figures 7-8; Supplementary Figure S5). The sweep on chromosome 15 occurred within Segment A (Figure 3) and showed no signs of introgression in any samples.

Several genes within sweep regions have potential roles in immune response and pesticide resistance (Supplementary Table S4). The sweep on chromosome 15 (Figure 7) contained six unique gene annotations, with the vast majority of CLR outliers observed within one gene – *Hyd*-like – encoding an immune-associated E3 ubiquitin-protein ligase (Cammarata-Mouchtouris et al. 2020). The sweep on chromosome 25 contained 7 functional annotations (Figure 6) and the sweep on chromosome 13, contained two genes including a different cytochrome P450 gene, *CYP333B3* (Supplementary Table 3; Figure 8).

### Evidence of strong, recent selection at *CYP333B3*

The largest sweep occurred at the cytochrome P450 gene *CYP333B3* on chromosome 13. This gene is of interest because of its validated functional association with pesticide resistance in *Helicoverpa*, and because nonsynonymous substitutions in this gene are known to have increased in frequency in North American *H. zea.* Assuming direct selection at this locus, we estimate the selection coefficient *ŝ* = 0.0489 (Figure 8A). This estimate is based on estimates of the effective population size, the recombination rate, and qualities of the sweep (see Methods for discussion of our simplifying assumptions and sources of error).

In North American *H. zea*, Taylor *et al*. (2021) identified temporal genetic differentiation concentrated in a region that tightly overlapped with the sweep we observed on chromosome 13. In the same study, the authors were able to go back to *H. zea* samples from 1998, 2002 and 2017 in their freezer collection to genotype individuals at this locus, showing that the proportion of individuals with derived non-synonymous *CYP333B3* mutations had increased over time. This afforded us an opportunity to determine whether our estimate of the selection coefficient was consistent with the change in allele frequency that they observed.

Our estimate of *ŝ* predicted the allele frequencies observed by Taylor *et al*. with less than 3% error when we assumed a dominant mode of inheritance and a biologically realistic generation time of 8-10 generations per year (Figure 8A, Supplementary Table S6). We also fit an estimate of the selection coefficient (*ŝ_fit_*) to the allele frequencies empirically estimated by Taylor *et al*. and found that the two selection coefficient estimates differed by less than 0.05 for generation times between 5 and 10 generations/yr (Supplementary Table S6). The dominance coefficient that best explained the data was >0.997 regardless of generation time when jointly optimised with the selection coefficient, consistent with our assumption of complete dominance. Therefore, we were able to precisely approximate the strength of selection at this locus from a single-timepoint sample. Moreover, this approximation was accurate enough to infer that anthropogenic selection led to a fixation of a derived *CYP333B3* allele within ∼20 years.

Patterns of genetic differentiation and diversity confirm that the selected allele arose within *H. zea* and was subject to selection within the last 20 years. When comparing *H. zea* samples from 2002 and 2019, genetic differentiation is greatest at the *CYP333B3* locus and otherwise low, consistent with a selective sweep occurring primarily in the intervening time (Supplementary Figure S6A). By contrast, differentiation is high across the full extent of chromosome 13 when comparing 2019 *H. zea* samples to *H. armigera* – especially at the *CYP333B3* locus (Supplementary Figure S6B). Among the 2019 samples, the average rate of coalescence was elevated at the locus. Specifically, there was a dearth of genealogies coalescing >2*N_e_* generations in the past at *CYP333B3*, (Supplementary Figure S6C) consistent with the effects of a selective sweep. These results contradict expectations under introgression (adaptive or otherwise), which would leave deep coalescence events at loci abutting the selected gene (Setter et al. 2020). On chromosome 13, the average height of coalescent genealogies estimated within individuals was 2*N_e_* generations (standard error 1.5×10^−4^ *N_e_* generations), contrary to what we would expect if some sampled haplotypes originated in a divergent population/species (see Methods). Together, these results indicate a strong selective sweep at *CYP333B3* over the past 20 years, and rule out introgression of *CYP333B3* from *H. armigera*.

### Signatures of selection at candidate Bt-resistance genes

None of the candidate Bt resistance loci that we mapped occurred in sweep regions, however one candidate locus on chromosome 13 occurred within the upper 1^st^ percentile of all CLR values (Figure 9). This locus (chr13:4012595-4041462) includes *PIK3C2A* and *kinesin-12*-like (positioned within *PIK3C2A*, on the opposing strand; Supplementary Table S4). Benowitz *et al*. (2022) identified a premature stop codon in *kinesin-12*-*like* (hereafter, *kinesin*-12) as the primary candidate target of selection underlying Cry1Ac resistance in the field-derived GA-R strain of *H. zea*.

Therefore, we investigated whether this same mutation occurred in our wild-caught samples. Requiring 95% confidence, we could call the presence or absence of the premature stop codon in 100 of our 237 samples. Of these, 99 carried the susceptible-strain allele and 1 carried a nonsynonymous C>A transversion mutating glutamine to lysine. We also identified 9 non-singleton SNPs in the coding region of *kinesin*-12 that were both nonsynonymous and resulted in amino acids with different biochemical properties to that of the susceptible strain (Supplementary Table S5). The most common of these was a C>A transversion mutating threonine to lysine, which occurred at allele frequency *q* = 0.11 (N=96 genotyped individuals). Therefore, the premature stop codon is unlikely to have caused the putative sign of selection we observed, though it is plausible that other nonsynonymous mutations have a similar phenotypic effect in disrupting the function of Kinesin-related protein 12. It is also possible that unobserved indels, larger structural variants, or cis-regulatory elements could cause the putative signal of selection observed at this locus. We note that the highest CLR values in this region align with segments immediately flanking *kinesin-12* (Figure 9).

## Discussion

### Gene flow within and between *Helicoverpa* species in North America

Our first aim was to test for the presence of invasive *H. armigera* ancestry in the north American *H. zea* population. Based on its rapid spread in South America, high propagule pressure, and availability of suitable contiguous habitat, Kritikos considered it was “a matter of time” until *H. armigera* – or introgressed *H. armigera* ancestry, at least – establishes in North America (Kriticos et al. 2015). Despite this, to our knowledge there has been no reported evidence of *H. armigera* in Central America. There has only been one report of *H. armigera* captured in the US prior to 2019, based on the COI haplotype of three individuals captured in Florida in 2015, with no evidence of establishment (Tembrock et al. 2019). We present the first conclusive report of admixed individuals in mainland North America. Only two individuals we sampled were admixed, and both were sampled at the same site in Texas despite an absence of population structure. The admixed samples show nowhere near the extent of *H. armigera* allele sharing observed in Brazil (Figure 1A). We saw no evidence of *H. armigera* introgression in previously reported samples collected in 2012 or 2017, though the sampling effort was far reduced in those years compared to our 2019 survey. Overall, the results point to a small degree of recent interspecific gene flow concentrated at a pesticide resistance locus.

Haplotypes break down over successive generations through recombination and mutation (Pool & Nielsen 2009). The size and therefore the detectability of introgressed ancestry blocks therefore also decays over time. Admixture between *H. armigera* and *H. zea* in South America was apparently punctuated by a pulse of hybridization 60-100 generations ago, with declining rates of hybridization since (Valencia-Montoya et al. 2020). We concluded that the vast majority of *H. zea* samples were non-admixed. Since we do not expect ABBA and BABA pattern frequencies to be exactly equal, values of *f_d_* insignificantly greater than zero across several chromosomes (Figure 1B) are consistent with an absence of introgression. This does not rule out the possibility of rare, short *H. armigera* haplotypes segregating in the *H. zea* population. However, our method was sensitive enough to detect *H. armigera* introgression in a region spanning ∼1% of the genome in <1% of individuals, suggesting that it was sufficient for our aim of identifying the presence of introgression at levels that are meaningful for the management of this pest.

Our second aim was to characterise population structure within non-admixed North American *H. zea*. While the use of genomic data has revealed otherwise cryptic population structure in some species, including *H. armigera* in its native range, we show that previous reports of complete panmixia in *H. zea* are not simply due to a lack of power (Zhang et al. 2022). Our analysis based on millions of SNPs showed that individuals collected at different locations in Texas were, on average, as genetically differentiated from one another as they were from individuals collected in North Carolina. While remarkable, this result is consistent with our understanding of *H. zea* movement ecology (Jones et al. 2018; Lawton et al. 2022). Given that summer incursions at latitudes higher than our sampling range are consistently repopulated by southern populations in the range we sampled, *H. zea* is likely panmictic across North America. Fitness-enhancing alleles can therefore spread readily across space. This highlights the need for monitoring and management at the species level across the agroecosystems potentially affected by *Helicoverpa*. Moreover, the evolution of pesticide or Bt resistance in one crop or region has immediate consequences throughout North America, where there may be substantially different agricultural and pest management practices (Reisig & Kurtz 2018). This result highlights the intrinsic ecological relationship between management practices in neighbouring fields, regions, or states, and should emphasise the need for multilateral management solutions.

### Pesticide resistance through interspecific introgression

*CYP337B3* could have been introduced via two routes: northward dispersal of admixed individuals from South America, or an independent introduction of *H. armigera* from its native range. An independent introduction is highly unlikely as it would require many generations of hybridization and recombination. Yet we saw no evidence of *H. armigera* ancestry in 2012 and 2017. Gene tree reconstruction could not be used to identify a clear source population of the *CYP337B3* polymorphism due to a lack of phylogeographic signal within *H. armigera* (Figure 5). However, the very fact that *H. armigera* ancestry is concentrated around the *CYP337B3* locus, which is overrepresented among admixed individuals in Brazil due to selection, (Valencia-Montoya et al. 2020) strongly suggests that the polymorphism was introduced through northward dispersal.

Our results show that the admixed samples carried alleles that, due to their semidominant phenotypic effect, enable resistance to fenvalerate, deltamethrin and possibly cypermethrin (Rasool et al. 2014; Ni et al. 2023; Joußen et al. 2012). Thus, an otherwise absent resistance phenotype was introduced to the North American *H. zea* population via interspecific introgression. The introgression of this gene is an indirect result of adaptation to pyrethroid exposure both in the native range of *H. armigera* (Rasool et al. 2014; Ni et al. 2023; Joußen et al. 2012) and subsequently in South American *H. zea* (Valencia-Montoya et al. 2020; Durigan et al. 2017). As anthropogenic activity increasingly brings pest species into secondary contact, introgression could become an important mechanism of rapid adaptation. This is particularly relevant to pesticide resistance, which can in some cases be underlain by a modular genetic architecture. For example, adaptive introgression of a *vkorc1* allele from the western Mediterranean mouse *Mus spretus* into the house mouse *M. musculus domesticus* confers the latter with resistance to anticoagulant rodenticides (Song et al. 2011). Introgression of an *ARH* allele from Atlantic to Gulf killifish species enabled tolerance to anthropogenic pollutants, which was shown to be highly fitness-enhancing (Oziolor et al. 2019). Similarly, interspecific introgression in the fungal genus that causes Dutch elm disease, *Ophiostoma*, was associated with virulence (Hessenauer et al. 2020). In each of these cases, genetic variation already shaped toward a fitness optimum in one species was introduced to another at relatively high frequency – an adaptive process that can be substantially faster than selection on *de novo* mutations (Marques et al. 2019).

### Pesticide resistance through rapid adaptation in *H. zea*

Taylor *et al*. (2021) measured genetic differentiation in North American *H. zea* between 2002 and 2017, showing that a region overlapping with the sweep we identified on chromosome 13 was the most significant outlier. This result provides strong independent evidence that we have correctly identified a selective sweep. Two annotations lie within the sweep region: *carboxypeptidase Q*-like and the cytochrome P450 gene *CYP333B3*. *CPQ*-like is noteworthy because some carboxypeptidases bind Cry1Ac in *Helicoverpa armigera* (Da Silva et al. 2018) and are up-regulated upon Bt exposure in other lepidopteran pests (Yang et al. 2018; Van Munster et al. 2007), though to our knowledge have never been functionally associated with Bt resistance. Selection at *CYP333B3* is a more parsimonious explanation for the sweep. Cytochrome P450s have repeatedly been implicated in the evolution of resistance to a range of pesticides, especially in *Helicoverpa* (Nauen et al. 2022). The P450 gene *CYP333B3* plays a general role in xenobiotic metabolism in *Helicoverpa* and other noctuid crop pests (Amezian et al. 2021; Shi et al. 2022). Studies of noctuid pests have shown that the gene is induced by a broad array of pesticide classes: indoxacarb (a voltage-dependent sodium channel blocker), fluralaner (a GABA-gated chloride channel allosteric modulator), imidacloprid (a nAChR competitive modulator), aldrin (an organochloride), several host plant defences (xanthotoxin and gossypol), and the pyrethroid fenvalerate (Amezian et al. 2021; Shi et al. 2022; de la Paz Celorio-Mancera et al. 2011; Han Yangchun 2014). Two key observations link *CYP333B3* evolution to pesticide resistance in *Helicoverpa*. First, Shi *et al*. performed an in vitro metabolism assay in *H. armigera* and found that *CYP333B3* showed the highest activity for metabolism of the organochloride aldrin (Shi et al. 2022). Second, Han *et al*. showed that *CYP333B3* is constitutively overexpressed in the pyrethroid strain FenR (Han Yangchun 2014). *CYP333B3* is therefore a key gene of interest with respect to pesticide resistance.

We estimated that *CYP333B3* was subject to a selection coefficient *ŝ* = 0.049 and showed that this coefficient could explain independently observed changes in the frequency of nonsynonymous mutations from 0.07 in 1998 to 0.87 in 2017 (Taylor et al. 2021). *H. zea* resistance to pyrethroids was first reported in 1990, and control failures became apparent in the South and Mid-West in the mid-2000s when derived *CYP333B3* allele frequencies were approaching 0.4 (Jacobson et al. 2009). By the late 2010s, pyrethroids were considered to have mixed efficacy compared to more effective diamide pesticides (Olmstead et al. 2016). So although further functional studies are needed to confirm the mechanistic link between the derived *CYP333B3* and pyrethroid resistance, two key observations lead us to hypothesise that pyrethroid exposure has caused rapid adaptation at this locus over the past two decades. First, there is clear evidence of a selective sweep at *CYP333B3* over the same narrow window of time in which phenotypic change was observed in pyrethroid resistance in North American *H. zea*. Second, there is a has been a detailed functional association between *CYP333B3* (its wild type genotype in *H. armigera*, at least (Han Yangchun 2014)) and those same pyrethroids. Regardless of the specific pesticide(s) that imposed positive selection on derived *CYP333B3* alleles, it is clear that the adaptive response occurred within a ∼20 year time frame – an example of rapid anthropogenic adaptation. Although *CYP333B3* appears to be under convergent selection in *H. armigera* in Europe, Africa and South America (Jin et al. 2023), we see no evidence of introgression into *H. zea* from *H. armigera* at this locus. Rather, the increased rate of coalescence and genetic differentiation from *H. armigera* and at this locus, combined with the recorded increase in frequency from a rare variant in *H. zea* over a decade before the first observation of *H. armigera* in South America, strongly supports selection from *de novo* mutation within *H. zea*. Thus, while *de novo* adaptation may be slower than adaptive introgression, it can occur at timescales once thought to be impossible for an evolutionary process. Therefore, for species with large and highly connected populations exposed to strong selection pressures, management strategies must not only consider ecological impacts but also evolutionary change over the short- and medium-term.

### Other signatures of selection in North American *H. zea*

Bt resistance in *H. zea* is a major concern. Extensive research has shown that resistance to some toxins (*e.g.* Cry1) is already common, and that selection for resistance to Cry2Ab2 was ongoing at the time we collected samples (Bilbo et al. 2019; Ali et al. 2006; Yu et al. 2021; Reisig et al. 2018; Dively et al. 2016). Although phenotypic data clearly show that resistance is evolving, identifying its genetic basis in both the field and the lab has been challenging. This is because the same phenotype can result from many polygenic, semi-overlapping genetic architectures. Multiple recent studies have identified Bt resistance QTL that do not overlap with any established Bt resistance loci (Taylor et al. 2021; Benowitz et al. 2022). Detecting the signature of selection on such complex traits is difficult without extensive replication or time-series data. Taylor *et al*. (2021) identified several QTL associated with Bt resistance but found no evidence of allele frequency shifts at these loci over time. It is perhaps not surprising, then, to find that none of the 19 Bt resistance QTL we investigated stood out as showing clear selective sweeps. One notable exception is the locus *PIK3C2A*/*kinesin-12* (Figure 9), in the upper first percentile of CLR values genome-wide. Benowitz *et al*. (2022) identified a premature stop codon in *kinesin-12* as the primary Bt resistance candidate in their QTL. We did not observe this mutation in our field-collected samples, though we did see non-singleton, nonsynonymous mutations in the coding region of *kinesin*-12 that altered the physio-chemical properties of amino acids and may have had the same phenotypic effect on disabling protein function. Cis-regulatory mutations could also disable the function of Kinesin-related protein 12; consistent with this hypothesis, the highest CLR values abut *kinesin*-12. So while the specific mutation observed by Benowitz *et al*. likely does not explain the putative signature of selection we observe, it is plausible that there has been selection at this locus in the field. Two important caveats should be noted. First, we could confidently call genotypes in ∼40% of our samples, so we cannot rule out selection on the premature stop codon itself in the field. Second, this locus is only noteworthy because of our specific candidate gene search; it would not have been included amongst the most likely targets of interest in our selective sweep analysis alone. Even though it is within the upper first percentile of CLR outliers, CLR values are distributed such that hundreds of other genes show more obvious signatures of selection (Figure S5). Therefore, functional work is needed to test Benowitz’s hypothesis that a premature stop codon in *kinesin-12*-like contributes to Bt resistance in the field.

To define sweeps in our hypothesis-free selection scan, we used a stringent yet arbitrary (upper 0.01^st^ percentile of CLR values) in order to avoid false positives, though this means that real selective sweeps may be missed. For example, chromosomes 10 and 26 showed CLR values only 4 units below the threshold (Figure S5). For two of the three putative sweeps we identified, we can only speculate on possible causes of selection. The sweep on chromosome 15 includes a *takeout*-like gene. *Takeout*-like genes are over-expressed in pyrethroid resistant mosquitos and aphids; in the latter RNAi experiments showed that they directly contribute to resistance (Peng et al. 2021; Toé et al. 2015).

*Takeout-*like genes were also significantly over-expressed in *Spodoptera litura* upon exposure to the isoxazoline insecticide fluralaner, and in honeybees exposed to the herbicide atrazine (Jia et al. 2020; Wang et al. 2023). It is therefore possible that this sweep is due to selection on a cis-regulator of *takeout*-like, though showing this requires further investigation. It is also noteworthy that the chromosome 15 sweep overlaps with two genes encoding E3 ubiquitin-protein ligases involved in protein degradation and cell cycle regulation (*Hyd-*like and *RNF168*-like) (Flack et al. 2017). Both *Hyd* and *RNF68* have been implicated in host-virus protein-protein interactions, consistent with selection imposed by a viral outbreak (Lilley et al. 2010). Some of the clearest signatures of selection result from outbreaks of infectious disease (Obbard et al. 2011). This may be particularly likely in North American *H. zea*, where nucleopolyhedroviruses are often used as a means of biocontrol alongside Bt toxins and synthetic pesticides (Niedermann et al. 2017).

The cause of the putative sweep on chromosome 25 is less clear, though we note that the dynein gene in this region was found to be the most up-regulated gene in an RNA-seq comparison of neonicotinoid-susceptible vs. resistant honeybees (Bahia 2021). While there is insufficient *a priori* information to go beyond a simple description of the sweeps on chromosomes 15 and 25, the plausible cause of selection on chromosome 13 is more obvious.

### Pleiotropy, epistasis, and the evolutionary fate of *CYP337B3* and *CYP333B3* alleles

We identified two cytochrome P450 genes recently introduced to the *H. zea* metapopulation via completely distinct evolutionary processes. We have argued that both are associated with pyrethroid resistance. This begs the question: is there an epistatic interaction between *CYP333B3* and *CYP337B3*, and if so, what does this mean for the future spread of *H. armigera* ancestry? If the derived allele of *CYP333B3* is common and *CYP337B3* is rare, is the marginal fitness of the latter completely diluted in this population? Several factors need to be considered. First, resistance to a specific pesticide is a quantitative trait for which distinct cytochrome P450 genes often have additive effects (Brun-Barale et al. 2010; Tchouakui et al. 2021). Second, pyrethroids represent a vast and diverse set of pesticides. No single mutation confers complete resistance to all pyrethroids, and different mutations associated with pyrethroid resistance almost always contribute to resistance in a nonoverlapping set of pesticides. This can be the case for completely different classes of pesticide. In other words, both resistance alleles have pleiotropic phenotypic effects. Third, the derived *CYP333B3* allele was already common in North America when *CYP337B3* was introduced to South American *H. zea* populations in 2013 (Figure 8B). *H. zea* populations are well connected between North and South America, as evidenced by the spillover of *H. armigera* ancestry that we report here. Pyrethroids have been commonly used in both Brazil and the US for pest control in cotton and maize (Walsh et al. 2022). Given that *CYP337B3* was highly fitness-enhancing in South America, and given that it would likely have often occurred in a genomic background with this same *CYP333B3* allele, we have little reason to think it would not spread in North America as well. That said, a fourth factor to consider is demography. Due in part to the phase of the El Niño southern oscillation, a population expansion of *H. armigera* in Brazil shortly after its detection appears to have resulted in demographic swamping, producing a pulse of hybridization (Specht et al. 2021; Valencia-Montoya et al. 2020). By contrast, in 2019 we see a ‘trickle’ of *H. armigera* ancestry into North American *H. zea*. Even alleles under strong selection can be lost through drift at sufficiently low frequency. Therefore, the ongoing influx of *H. armigera* ancestry and the North American selective regime with respect to *CYP337B3* will together determine spread. While a degree of predictive power could be gained through fitness assays to determine the epistatic interaction between *CYP337B3* and *CYP333B3* for tolerance to fenvalerate and other commonly used pesticides, the outcome of this incursion remains to be seen through future biosurveillance.

### Prospects for genomic surveillance in this system and others

Early detection of invasive pests can pay dividends even when surveillance is costly, as invasive species are easier to manage shortly after they establish (Mehta et al. 2007). Genomic detection of *H. armigera* is currently far more scalable and sensitive as a diagnostic test to distinguish *H. zea* from *H. armigera* relative to morphological or single-locus genetic assays. Nonetheless, a trade-off exists between the cost of surveillance and the product of the probability and cost of invasion. Given that *H. armigera* ancestry has spread into the United States, and the value of the crops exposed to *H. armigera* (∼US$78 billion per annum (Kriticos et al. 2015)), there is a strong case for extensive ongoing genomic monitoring of *H. zea* across North America, especially in the dispersal corridor leading through Central America and Mexico where there has been comparably little genomic monitoring to our knowledge. Monitoring programs must adopt a genic view of biological invasion in order to quantify the incursion of *H. armigera* (North et al. 2021). This means that, if the aim is to map the geographic distribution of possible *H. armigera* spread, *CYP337B3* assays are likely to be more informative than ancestry-based assays that use one or few markers. Alternatively, if the aim is to determine the presence of *H. armigera* introgression in an area of concern, only genomic data offer the resolution required to detect the short haplotypes segregating in the population. Future work to characterise the sets of loci most sensitive to admixture (*e.g.* those in high-recombining regions, least likely to be removed by selection due to linkage with Bateson-Dobzhansky-Muller incompatibilities) will allow for the development of SNP arrays that can robustly assign ancestry proportions to samples at lower cost.

When genomic data was first adopted in the field of molecular population genetics, there was excitement about the power of population genomics to infer the evolutionary history and genetic basis of adaptive traits in wild populations, especially where laboratory crosses were impossible (Li et al. 2008). While genomic surveillance has become a critical technology for the management of pests and invasive species (Hamelin & Roe 2020; Chown et al. 2015), in practice the detection of adaptation can be limited by the complex genetic architectures and demographic nonequilibrium. Incorporating QTL metadata and sampling over a time-series can address these issues – reinforcing the need for sustained genomic monitoring programmes – though this is not always possible (Taylor et al. 2021; Clark et al. 2023; Pélissié et al. 2018). Although we only sample individuals at a single timepoint, we show that relevant population genetic statistics can be estimated with sufficient accuracy to not only identify a putative target of selection for pesticide resistance, but to precisely infer recent allele frequency change at the locus. The analyses we apply here demonstrate the richness of information that can be extracted from nucleotide diversity in wild pest populations to inform management action and study anthropogenic adaptation.

In summary, we have shown that pesticide resistance in North American *H. zea* arose recently, and rapidly, via two independent processes: interspecific introgression and intraspecific adaptation. These findings underscore the importance of rapid adaptation for pest management – evolution at timescales previously only considered to be relevant for ecological process.

## Methods

### Sampling, DNA extraction, library preparation and sequencing

*H. zea* individuals were collected in agricultural fields at 10 locations across the southern U.S. in 2019 (see Supplementary Table S1 and Figure 1) using corn earworm pheromone lure (GreatLakes IPM, Vestaburg, MI). Adult moths were immediately placed in 95% ethanol and stored at −20°C prior to DNA extraction. A few populations of *H. zea* were collected as larvae (see Supplementary Table S1) and brought back to the laboratory of Dr. David Kerns at the Texas A&M University Dept. of Entomology in College Station, TX, USA where they were raised to adults and then processed for DNA extractions.

The abdomen of each adult moth was separated from the body and used for DNA extraction with the Qiagen DNeasy Blood & Tissue kit (Qiagen, Germantown, MD following the manufacture’s protocol. The final DNA elution step was performed using Qiagen buffer EB (Qiagen, Germantown, MD) instead of the AE buffer. DNA concentration was measured using a Qubit 3.0 Fluorometer (Thermo Fisher Scientific, Waltham, MA). Genomic DNA samples were delivered to the Texas A&M AgriLife Genomics and Bioinformatics Service in College Station, TX for library preparation and sequencing.

DNA batch purity and integrity were assessed using a DS-11 spectrophotometer (Denovix, Wilmington, DE) and capillary gel electrophoresis (Fragment Analyzer, formerly Advanced Analytical, now Agilent, Santa Clara, CA), respectively, from 11 samples per 96-well plate. Genomic DNA was further purified with SPRI beads (Omega Bio-tek, Norcross, GA) and the concentration of each sample was determined using Lunatic plates (Unchained Labs, Pleasanton, CA) read on a DropletQuant spectrophotometer (PerkinElmer, Waltham, MA).

Libraries were prepared from 33ng of genomic DNA using a custom miniaturized version of the NEXTFLEX Rapid XP kit protocol (PerkinElmer) that was automated on a Sciclone NGSx liquid handler (PerkinElmer). Briefly, all reactions involving enzymes were carried out in one third of manufacturer proscribed volumes for each reagent. In the modified reaction volumes, genomic DNA was enzymatically fragmented for 3 minutes, then ligated to unique dual-indexed barcodes. The raw library reaction was brought up to protocol volumes with H_2_O for a SPRI cleanup followed by SPRI size selection between 520 and 720 bp, and then eluted in reduced volume to be amplified for 10 PCR cycles. Finally, the amplification reaction was brought back up to recommended volume with H_2_O to perform an additional single sided SPRI size selection that retains DNA fragments larger than 450bp.

A subset of libraries was checked for size and integrity on the Fragment Analyzer and all libraries were quantified using a fluorescent plate reader (SpectraMax M2, Molecular Devices, San Jose, CA) with PicoGreen reagent as per the manufactures suggested protocol (Thermo Fisher Scientific, Waltham, MA). Libraries were diluted with EB (Omega Bio-tek) to a final concentration of 2.25 ng/µl using an automated liquid handler (Janus, PerkinElmer) and an equal volume of each was pooled. Pooled library quality was assessed on the Fragment Analyzer and molarity determined using a qPCR-based Library Quantification assay (Roche, Pleasanton, CA).

The pool was sequenced in a single lane of an Illumina NovaSeq S4 XP flowcell (San Diego, CA) using the 2 × 150 bp recipe. The raw data was demultiplexed with bcl2fastq 2.20, which yielded 2.17 billion demultiplexed reads ranging from 5.01-10.41M reads per sample and an average of 7.80M reads per sample (∼6.5x coverage).

### Filtering, alignment, and variant calling

In order to conduct intra- and inter-specific comparisons, we used raw sequencing data from multiple *Helicoverpa* spp. generated by Anderson *et al*. (2018), Taylor *et al*. (2021) and Jin *et al*. (2023). These publicly available data were analysed in the same manner as our newly generated sequence data. Two slightly different pipelines were used for different analyses depending on the samples required. Bioinformatics pipelines were implemented using Snakemake v7.2 (Mölder et al. 2021).

For the results presented in Figures 1-6 (hereafter, Call Set 1): Fastq files were trimmed using fastp v0.23 (Chen et al. 2018), then mapped using bwa v 0.7.12 (Li & Durbin 2009). The resulting .bam files were sorted using SAMtools (Li et al. 2009). Picard v2.9.2 (Broad Institute 2023) was used to remove duplicates and SAMtools was used to index the filtered .bam files. GATK HaplotypeCaller v4.3 (van der Auwera et al. 2013) was used to call haplotypes per individual. GATK CombineVCFs was used to merge call sets across all individuals and GenotypeGVCFs was used to jointly call genotypes across samples sequenced here, those produced by Anderson *et al*. and samples collected in 2002 by Taylor *et al*. (2021). Of these, only single nucleotide polymorphisms were retained. VCFtools (Danecek et al. 2011) was used to filter out sites with a phred-scaled quality score below 20, sites with a mean depth of coverage below 2X or above 200X, genotypes with a mean depth below 2X or above 200X, with a missing data threshold of 50%.

**Figure 1:**
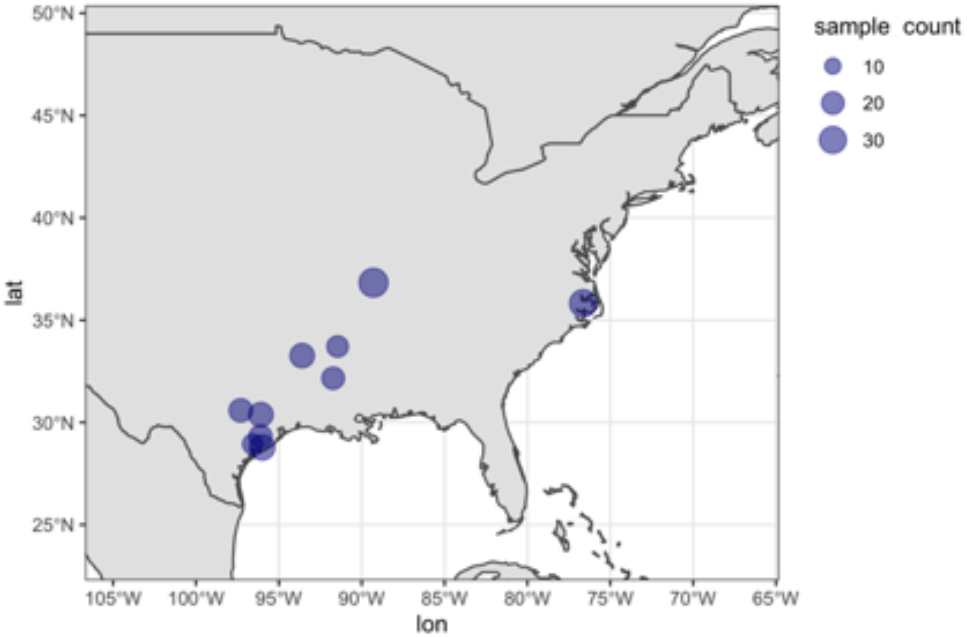
Individual sampling effort at 10 sites in 2019. Sample site information is detailed in Supplementary Table S1.

**Figure 2:**
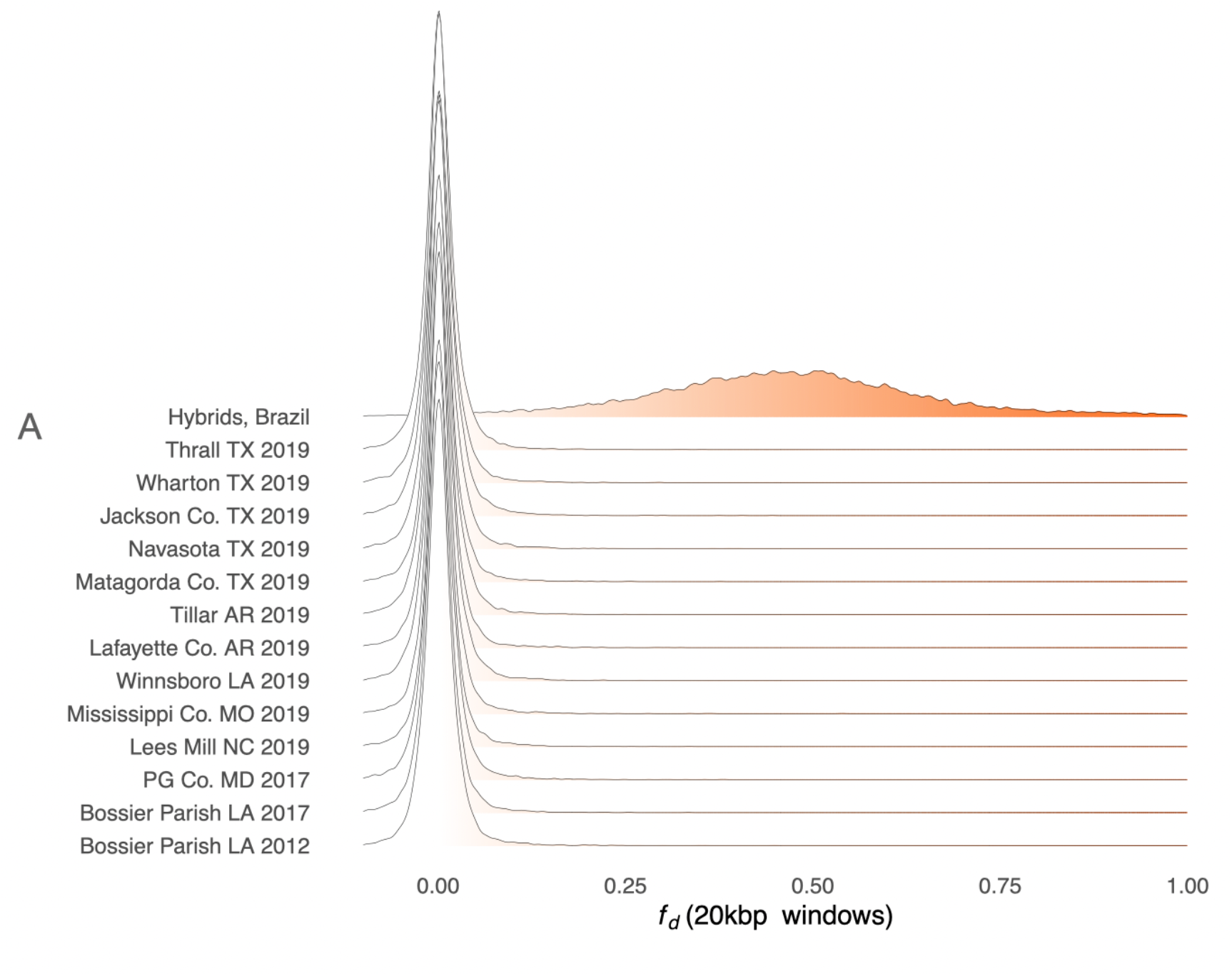
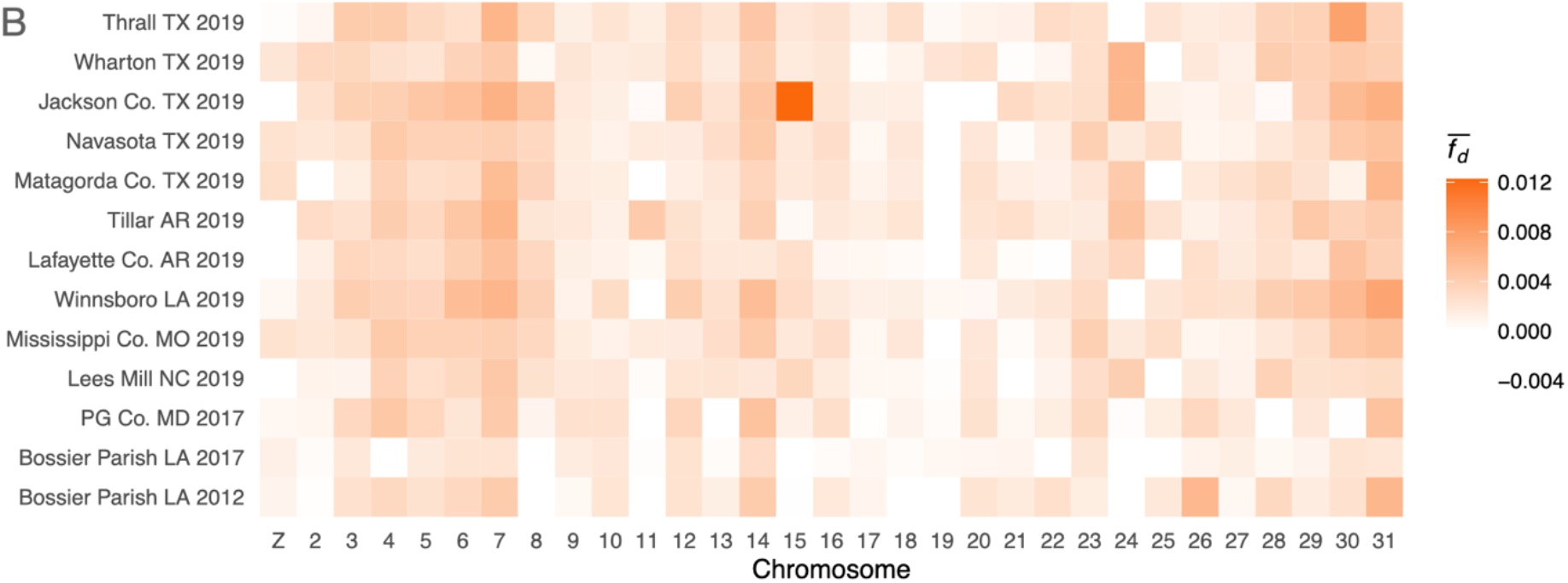
No evidence of *H. armigera* in *H. zea* except for on chromosome 15 among a subset of *H. zea* individuals collected in Jackson County, TX, in 2019. A: Distribution of 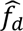 calculated in 20kbp windows where P1: *H. zea* sampled in 2002, P3: *H. armigera*, outgroup: *H. punctigera*. The statistic was calculated for 14 different P2 sets: 10 sets of *H. zea* samples collected at different sites in 2019, 2 sets of *H. zea* collected in 2017, one set of *H. zea* collected in 2012, and a positive control of 9 individuals sampled in Brazil shown to be admixed individuals carrying both *H. armigera* and *H. zea* ancestry. B: Data presented as the mean per chromosome.

**Figure 3:**
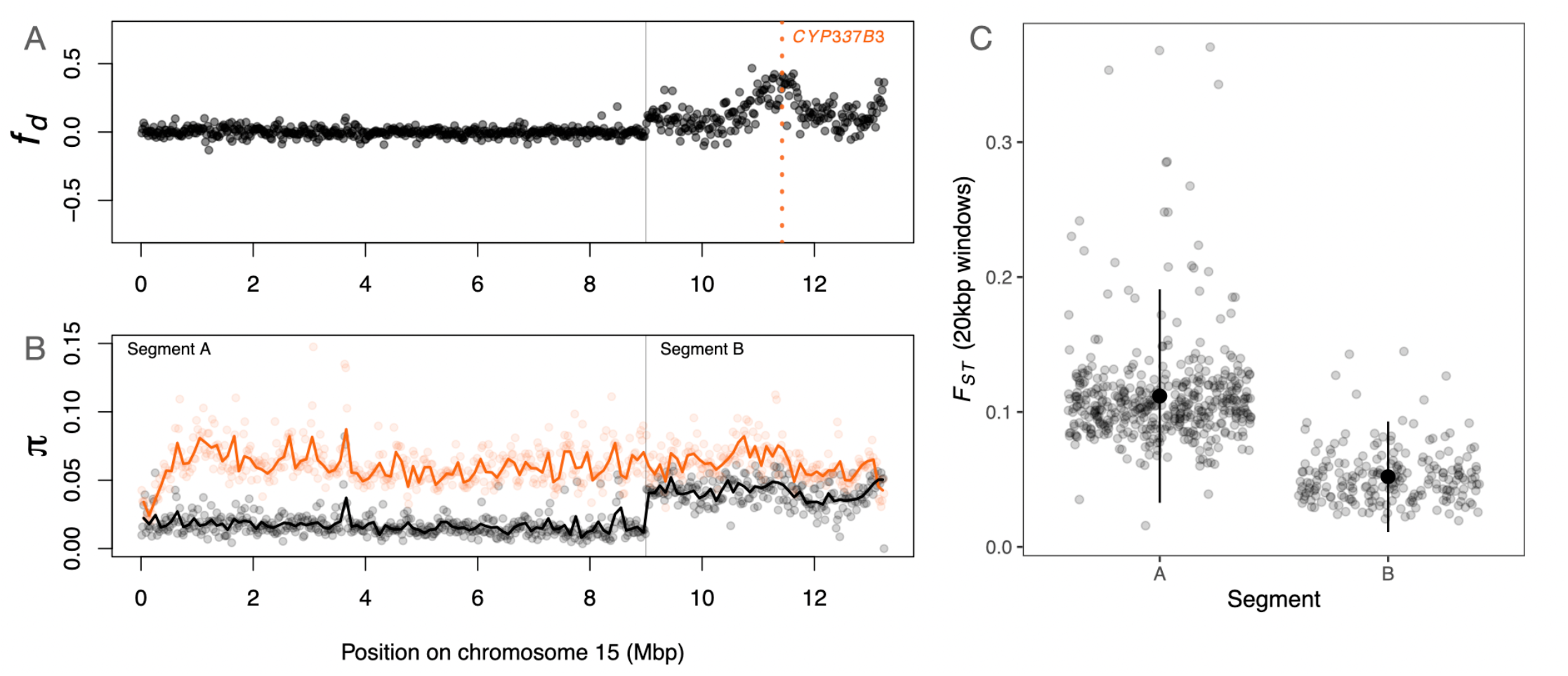
Introgression of *CPY337B3*. A: 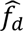 calculated in 20kbp windows along chromosome 15, where P1 is *H. zea* sampled in 2002, P2 are Ja15 and Ja25 in supplementary Figure S1, P3 is *H. armigera*, and the outgroup is *H. punctigera*. B: Nucleotide diversity (*π*) calculated in 20kbp windows (points) and 100kbp windows (lines) for the admixed individuals (Ja15 and Ja23; black) and for *H. armigera* (orange). C: Genetic differentiation (*F_ST_*), calculated in 20kbp windows, between the admixed individuals and *H. armigera* in the chromosomal segments labelled in (B).

**Figure 4:**
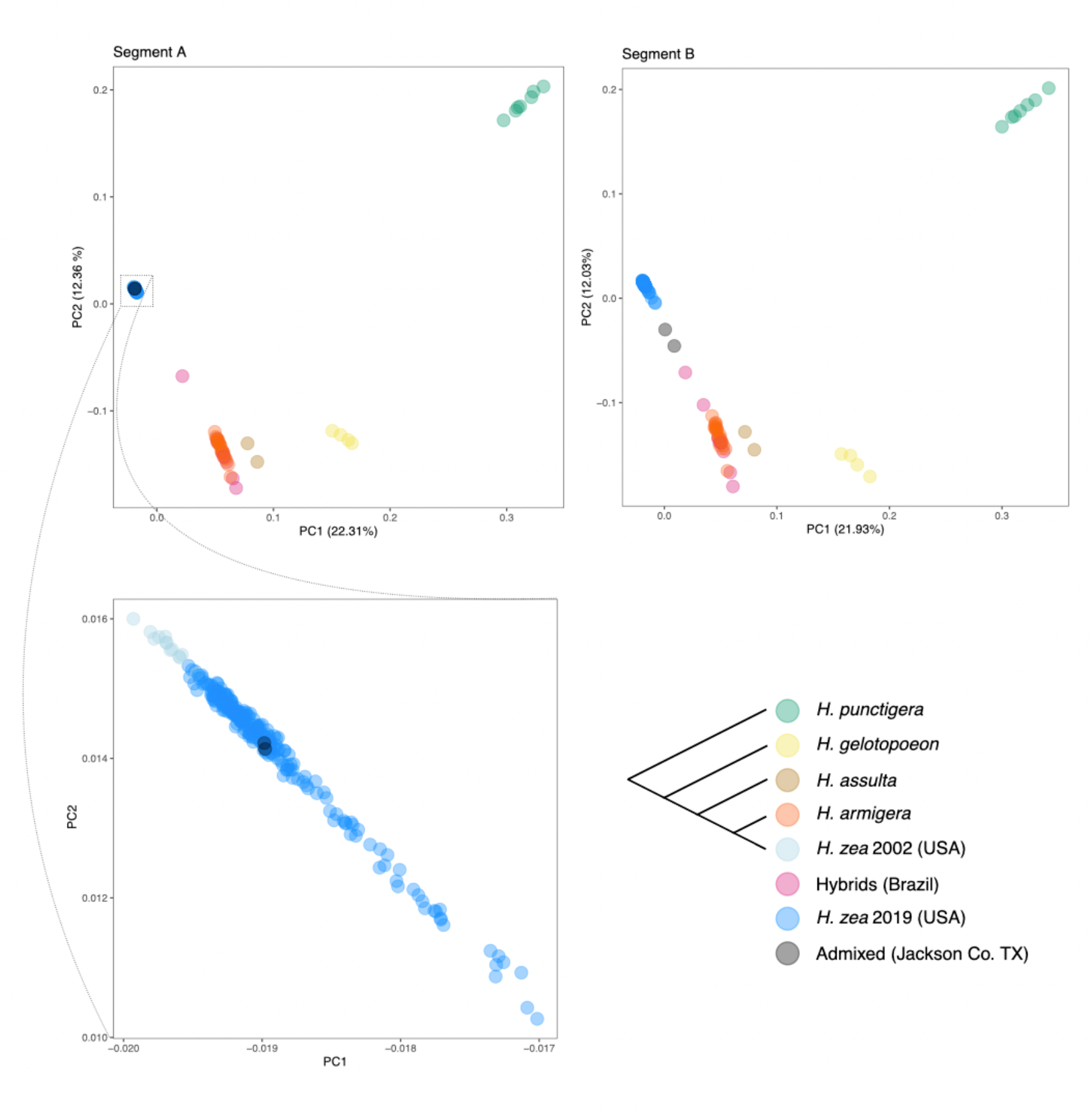
Visualising local ancestry with Principal Components Analysis. Principal components 1 and 2 calculated using SNPs form segments A and B of chromosome 15 (see Figure 3). Phylogeny in bottom right panel indicates the consensus species tree. In segment A, the two admixed samples (black points) cluster with other *H. zea* samples collected in 2019. In Segment B, the two admixed samples move in PC space toward *H. armigera* samples and admixed samples from Brazil.

**Figure 5:**
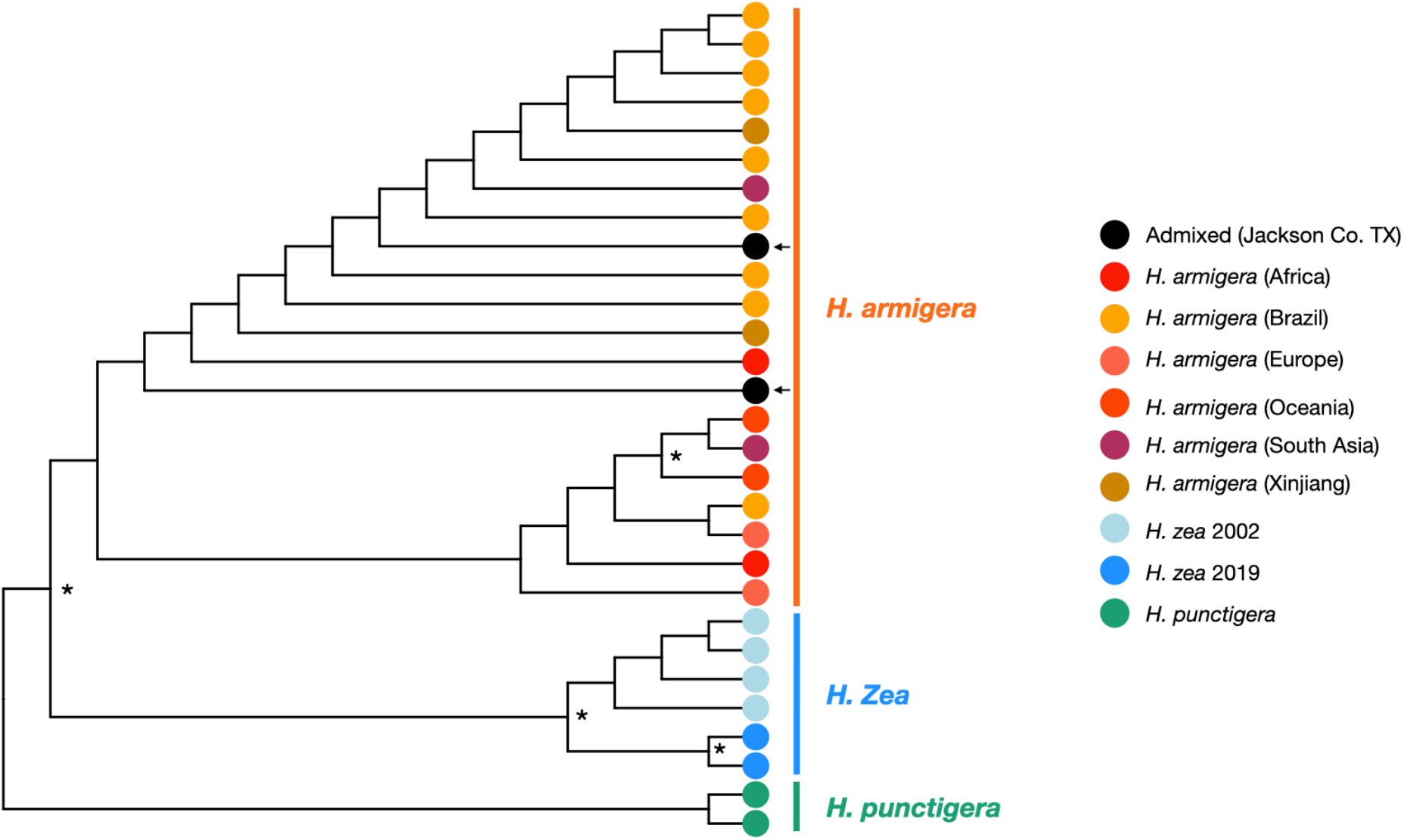
*At the CYP337B3* locus, *admixed samples form a clade with H. armigera samples*. Maximum likelihood cladogram rooted with *H. punctigera* samples. Asterixis denote nodes with >95% bootstrap support. Arrows indicate admixed samples. Horizontal lines group species-level clade. There was consistent bootstrap support separating species-level clades and *H. zea* samples from 2002 versus 2019, but not within *H. armigera*.

**Figure 6:**
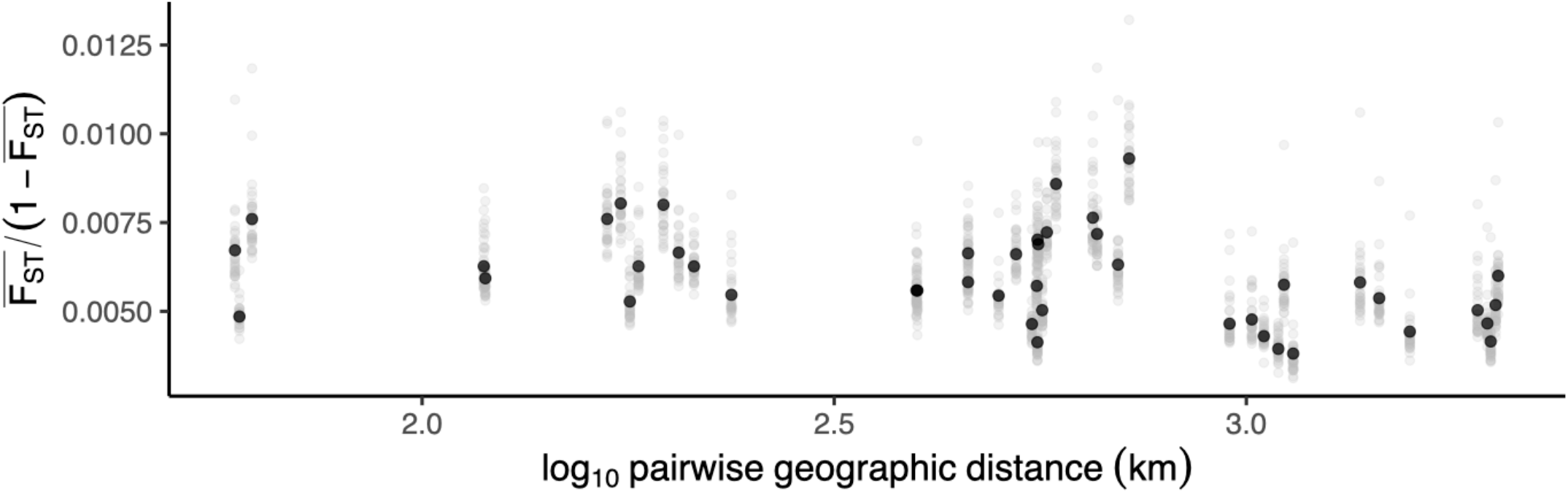
Low genetic differentiation irrespective of geographic distance among non-admixed H. zea. Pairwise genetic differentiation regressed on pairwise geographic distance for each of 45 possible comparisons among 10 sample sites. There is no positive relationship. Grey points are results for each of 31 chromosomes; black points are genome-wide averages. Genetic differentiation was calculated as mean *F_ST_* across 20kbp windows.

These samples were mapped against the highly contiguous *H. armigera* reference genome to maximise sensitivity when detecting introgression from *H. armigera*, and to allow direct comparison with results generated in similar studies (Taylor et al. 2021; Valencia-Montoya et al. 2020). The *H. armigera* genome is highly contiguous with the *H.* zea chromosome except for the sex chromosome (Benowitz et al. 2022). Therefore, some *H. zea* sex chromosome or pseudo-autosomal reads miss-mapped to autosomal scaffolds in the reference assembly. To avoid potential biases introduced through miss-mapping, Plink v1.9 (Chang et al. 2015) was used to calculate the statistical association between each autosomal SNP and heterozygosity on the Z chromosome, in which Z chromosome heterozygosity was treated as a quantitative trait. SNPs with *p* values in the top first percentile of association were excluded from all downstream analyses. This resulted in a call set of 45,700,390 SNPs among 304 individuals of four species (Supplementary Table S2). For analyses requiring additional samples (*e.g. H. zea* from 2017 and 2012, and those shown in Figure 5) mapping and variant calling was completed a subset of relevant samples was jointly called using the same pipeline.

For the selection analysis (results presented in Figures 7-9; hereafter Call Set 2) a more stringent filtering pipeline was used. Paired end raw reads were trimmed using Trimmomatic v0.38 (parameters: LEADING:3 TRAILING:3 SLIDINGWINDOW:4:15 MINLEN:50) (Bolger et al. 2014). Trimmed and paired reads were aligned to the *H. armigera* reference assembly generated by Pearce *et al*. (Pearce et al. 2017). Alignment was performed using bwa (Li & Durbin 2009) and mate pairs were fixed using SAMtools v1.9 (Li et al. 2009). Resulting bam files were sorted using SAMtools. Picard v2.18 was used to add read groups, clean bam files, and mark duplicates. Subsequently, SAMtools was used to index the bam files, and GATK HaplotypeCaller v4.15 (van der Auwera et al. 2013) was used to call haplotypes individually for each sample. Haplotypes were imported into a database using GenomicsDBImport from GATK. Lastly, the module of GenotypeGVCFs from GATK was used to call SNPs from all samples jointly. Next, the called genotype in a vcf file was subjected to a sequential filtering process using VCFtools v0.1.16 (Danecek et al. 2011). First, we removed samples that had over 50% of loci missing; these samples usually resulted from poor sequencing coverage. Second, we applied a quality filter by removing loci that had a genotype quality score lower than 20 (minGQ 20). Third, we applied a depth filter to remove loci that had coverage less than 2 or greater than 200. We only kept biallelic SNPs and removed all indels. Next, we removed loci missing from more than 50% of the samples missing, as well as all singletons in the vcf file. In the next step, we filtered out loci that violated Hardy-Weinberg equilibrium (HWE). HWE filtering was only conducted for *H. armigera* and *H. zea* samples because of the much smaller sample size of other species. HWE tests were applied only to loci with no missing genotypes, and *p*-value cutoff was set to 0.01. Loci that violated HWE in two of the three species were removed. We imputed the missing genotypes using Beagle (v5.1, Browning et al.) with default settings. Then we only kept biallelic SNPs in the imputed genotypes, and we also applied linkage disequilibrium pruning using Plink (--indep-pairwise 50 10 0.5). SNPs associated with Z chromosome heterozygosity were filtered using the method described above, resulting in a set of 5,706,238 SNPs across 237 *H. zea* individuals.

**Figure 7:**
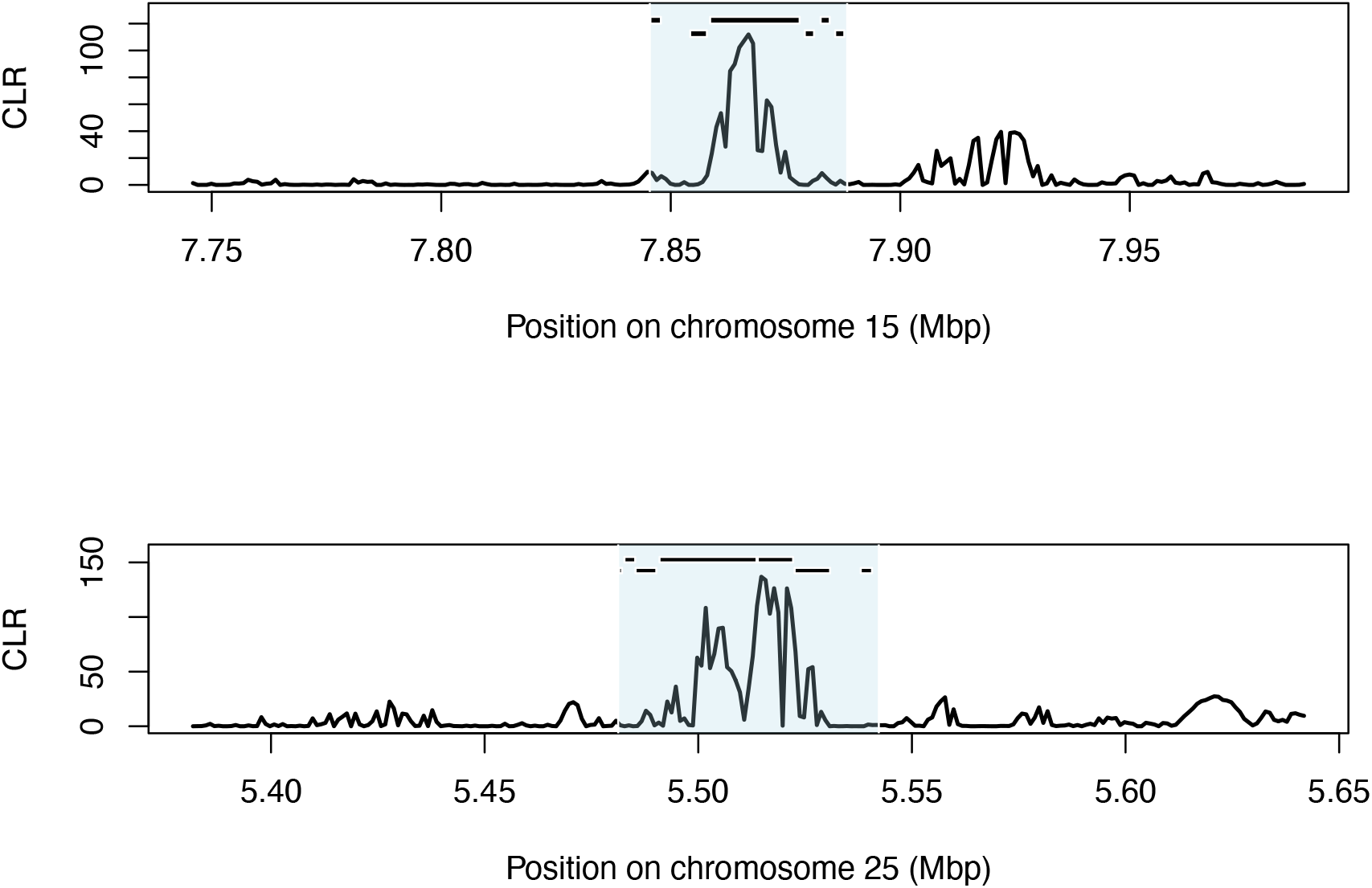
Sweep regions on chromosomes 15 and 25. Composite likelihood ratio (CLR) calculated in SweepFinder2. Black bars indicate the position of gene annotations on the forward strand (top) and reverse strand (bottom) listed in Supplementary Table S3. Blue shading indicates the sweep region.

**Figure 8:**
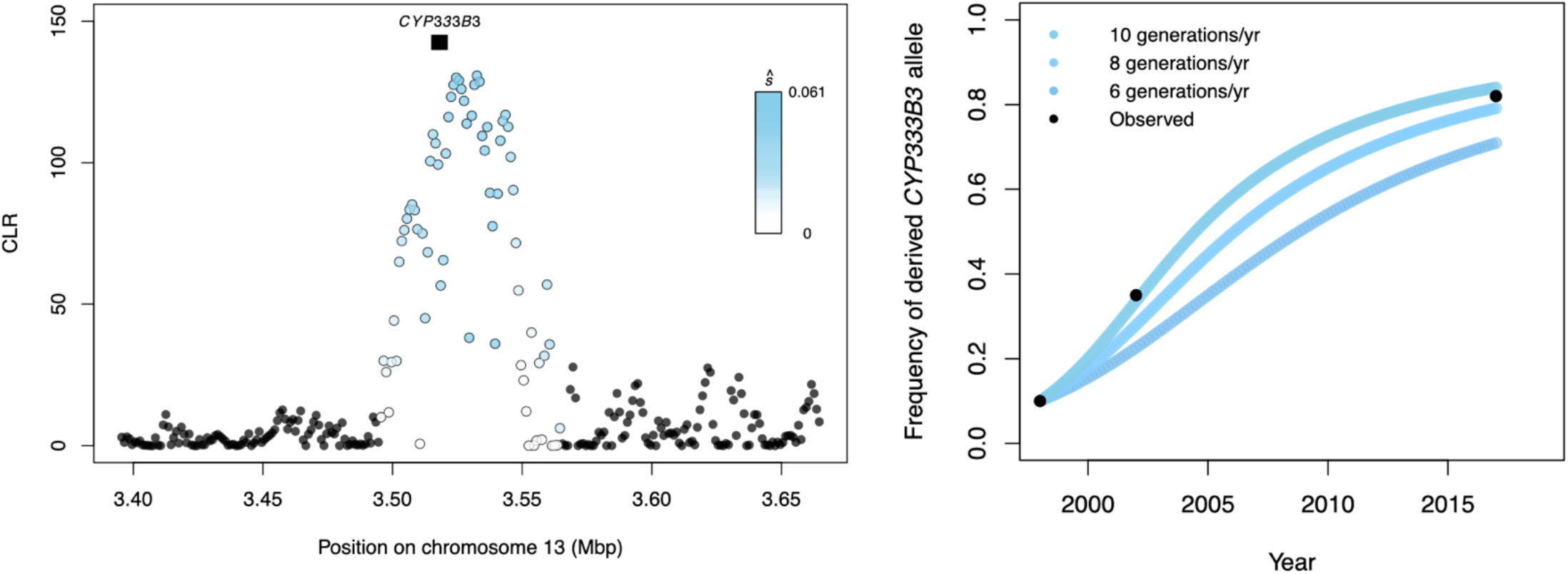
Selective sweep at *CYP333B3*. **A:** Composite likelihood ratio calculated in SweepFinder2, with points coloured by their estimated selection coefficient. Black box indicates the position of *CYP333B3*. **B:** Predicted frequency of a dominant-acting derived *CYP333B3* allele for each generation given the estimated selection coefficient at that locus *ŝ* = 0.0489 assuming 6, 8 and 10 generations per year. Point colour corresponds to assumed generation time. Black points are independently estimated allele frequencies from Taylor *et al*. (2021).

**Figure 9:**
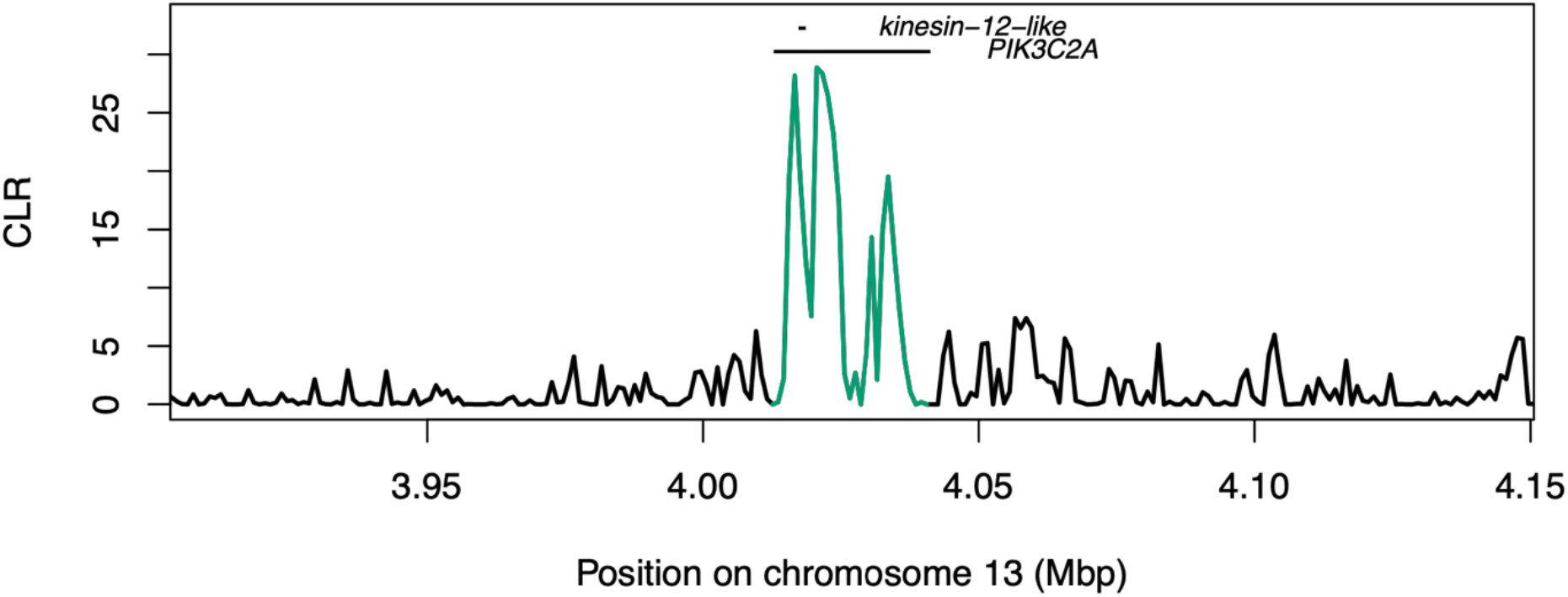
Selective sweep likelihood ratio at the locus *PIK3C2A/kinesin-12*-like. CLR values in the region of the loci *PIK3C2A* and *kinesin-12*-like, encoded on opposing strands. Both genes are candidate Cry1Ac resistance loci. Values overlapping with the annotation positions are highlighted in green.

For most window-based analyses described below, a window size of 20kbp was chosen because linkage disequilibrium decayed to a genome-wide background level of *r*^2^ = 0.002 over approximately 20kbp for the population of North American *H. zea* that we sampled.

### Detecting introgression

To test for allele sharing with South American *H. armigera,* we calculated 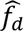 in 20kbp windows across all chromosomes in which at least 200 SNPs were called, using Python scripts described in Martin in *et al*. (Martin et al. 2015). P1 was designated as a set of 13 *H. zea* individuals sampled in 2002 in Louisiana by Taylor *et al*. (2021). These samples were collected over a decade before *H. armigera* was detected in the Americas and therefore represent the best possible whole-genome reference set of non-admixed *H. zea*. Note that we do not use *H. zea* individuals sampled in New York in 2005 from Anderson *et al*. (2018), as the metadata associated with these samples was contradictory and could not be verified. P3 was 25 non-admixed *H. armigera* sampled in Brazil, and the outgroup was 7 *H. punctigera* individuals sampled in Australia. For our samples, 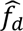 was calculated 10 times where the test set, P2, was *H. zea* sampled in 2019 at each of the 10 samples sites. And additional three tests were carried out *H. zea* collected in 2012 in Louisiana, and collected in 2017 in both Maryland and Louisiana by Taylor *et al*. (2021). As a positive control, 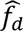 was also calculated for 9 individuals sampled in Brazil shown to be admixed offspring of *H. armigera* and *H. zea* by Anderson *et al*. (Anderson et al. 2018). Results from all 14 tests are shown in Figure 1.

### Calculation of summary statistics and visualisation with PCA

To compare the two admixed individuals with *H. armigera* samples we calculated nucleotide diversity (*π*), genetic differentiation (*F_ST_*), and genetic divergence (*d_xy_*) for the results presented in Figure 3 and Supplementary Figure S2 using python scripts described by Martin *et al*. (Martin et al. 2015). This was done in 20kbp and 100kbp windows (with at least 200 and 100 informative sites, respectively) with consistent results. Principal components analysis (Figure 4) was carried out using Plink v1.9 (Chang et al. 2015) in segments A and B separately.

### Reconstructing gene trees

To reconstruct a tree at the *CYP337B3* locus, we used a combination of *H. zea* samples collected here (two admixed and two representative non-admixed samples), 4 *H. zea* collected in 2002 by Taylor *et al*. (2021), 2 *H. punctigera* and 9 *H. armigera* collected from both the invasive range in Brazil by Anderson *et al*. (2018) (which are admixed with *H. zea*), and 10 *H. armigera* from the major clades of the native range by Jin *et al*. (2023) (see Supplementary Table S2). These samples were mapped, jointly genotyped and filtered without phasing in the manner as described above. We used VCFtools v0.1.15 (Danecek et al. 2011) to subset the *CYP337B3* locus previously used to identify signatures of adaptive introgression (HaChr15:11436565-11440168; Valencia-Montoya et al. 2020) from a multigenome VCF and converted to PHYLP format. We removed invariant and uninformative sites and retained sequences with no more than 50% missing data. We reconstructed a maximum likelihood phylogram the GTR+GAMMA model of rate heterogeneity and Lewis ascertainment bias correction method implemented in RAxML v 8.2.12 (Stamatakis 2014) with 100 bootstrap iterations. We rooted the tree using the two outgroup *H. punctigera* samples in ggtree (Xu et al. 2022), ensuring bootstrap values were assigned to the correct node with the edgelables function (Czech et al. 2017). The resulting cladogram is shown in Figure 5. The bootstrap values, tip label sample names and scaled branch lengths are shown as a phylogram in Supplementary Figure S4, which was rooted in FigTree v1.4.4.

### Measuring Isolation by Distance

To investigate connectivity across the North American *H. zea* metapopulation, we calculated the mean of *F_ST_* across 20kbp windows from all chromosomes for each of 45 possible pairwise combinations of the 10 sampling locations. Geographic distance was calculated for the same pairwise combinations using the R package geosphere (Karney 2013). We regressed 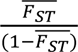 on transformed geographic distance to test for a positive correlation. A positive correlation would suggest a pattern of isolation-by-distance and provide a rough estimate of the product of population density and the variance in dispersal distance, whereas the absence of a correlation would indicate panmixia across sampling locations (Rousset 1997).

### Identifying selective sweeps

Briefly, we identified selective sweeps as localised and extreme deviations from the genome-wide site-frequency spectrum (SFS) consistent with linked selection using the composite likelihood ratio test implemented in *SweepFinder2* (DeGiorgio et al. 2016; Nielsen et al. 2005). To do so, we used sequence data from other *Helicoverpa* species to distinguish ancestral from derived alleles. By using the empirical SFS as a null model, rather than an equilibrium null SFS, we account for the confounding influence of demographic nonequilibrium (Nielsen et al. 2005). We use sequence data from other *Helicoverpa* species to estimate chromosome-wide mutation rates in order to estimate the population recombination parameter, then use chromosome-wide recombination rate estimates and estimates of the effective population size to estimate the strength of selection that acted at a locus of interest. Finally, we show that the estimated selection coefficient can explain independently observed shifts in allele frequency.

Specifically, we used VCFtools v0.1.15 (Danecek et al. 2011) to calculate allele frequencies within the North American *H. zea* population sampled in 2019. Including 25 non-admixed *H. armigera* individuals and 7 *H. punctigera* individuals as outgroups, we used a custom python script to designate ancestral and derived states to those alleles and calculate all possible transition and transversion rates for each chromosome. With these ancestral and derived states, *SweepFinder2* (DeGiorgio et al. 2016) was used to compute the unfolded genome-wide site frequency spectrum in the North American *H. zea* population, and to use this spectrum as a null model in a scan for selective sweeps. Briefly, *SweepFinder2* runs a composite likelihood ratio (CLR) test to identify chromosomal regions that show a decline in genetic diversity characteristic of genetic hitchhiking (Kim & Stephan 2000). The CLR was computed in 1kbp intervals. Note that the LD-based local recombination rate estimates generated below were not used to detect the location of selective sweeps, as this would have confounded the localised effect of selection on both the site-frequency spectrum and on linkage disequilibrium. We identified sites of interest (those clearly showing signs of selection) as the top 0.01^st^ percentile of CLR values (CLR>104.17). Sites above this threshold within 20kbp of one another were grouped into clusters using BEDtools v2.20.1 (Quinlan & Hall 2010), whereby the cluster start and end positions extended 20kbp up/downstream of the terminal outlier sites. We refer to these regions as putative selective sweeps.

### Mapping known candidate Bt resistance loci

We identified three classes of previously identified candidate Bt resistance loci: genes within the QTL described by Benowitz *et al*. (2022); genes repeatedly implicated in Bt resistance which were also examined by Taylor *et al*. (2021); and the novel resistance QTL identified by Taylor *et al*. (2021).

First, Benowitz *et al*. (2022) mapped a 250kbp Bt resistance QTL to chromosome 13 containing 10 genes of interest, including the putative causative locus *kinesin-12*-like, using a different reference assembly. To test for signatures of selection in this region, we used BLAST v2.4 to map each of the 10 loci in Table 2 of Benotitz *et al*. to our reference assembly, retaining the largest region of overlap with an *E*-value of 0 for each locus. As expected given the contiguity between our assemblies, all loci mapped within 250kbp of one another on chromosome 13. Within each BLAST hit, we used BEDtools v2.20.1 BEDops 2.4.41 to identify annotations in our assembly with matching annotation names (Quinlan & Hall 2010; Neph et al. 2012). Two annotations (*kinesin-12*-like, *JHE*-like) were absent in our annotation; we excluded the latter because we could not confirm its location. Because *kinesin-12*-like mapped well to our assembly (>90% sequence identity, *E*-value=0), and since *kinesin-12*-like sits within the larger gene *PIK3C2A* (where it is encoded on the opposite-sense strand) we could map the 1.7kbp gene with a range of error <50bp.

Second, we applied the same mapping approach to identify other known Bt resistance loci listed in Taylor *et al*. (2021) Table S9. We were able to confidently assign corresponding annotations for all genes except alp, cad2, *calp4* and *abcA2*.

Third, Taylor *et al*. (2021) identified many QTL associated with resistance to crops expressing both Cry1Ab and Cry1A.105 plus Cry2Ab2 toxins. Both traits were highly polygenic. We identified the most significant outlier scaffold for trait for both traits and mapped these to our assembly. This was done for separately for *H. zea* scaffolds and *H. armigera* super-scaffolds. To be consistent with Taylor *et al*. as much as possible, we defined the most significant outlier scaffold as that with the lowest mean *p* value for the linear mixed model likelihood ratio test with at least one SNP assigned a2 BSLMM posterior inclusion probability >0.01. We retained only super-scaffolds with at least 10 SNPs and scaffolds with at least 3 SNPs. We identified the scaffold NW_018395566 and the nested scaffold KZ118765 (nested within NW_018395566) as the scaffolds most associated with Cry1Ab resistance NW_018395399 and the nested KZ118015 were most associated with Cry1A.105+Cry2Ab2 resistance. All four were within the resistance-associated linkage group 9 in the map produced by Taylor *et al*. and neither were associated with growth rate on either of the control treatments. We mapped these scaffolds to our reference assembly, retaining the largest hit with an *E*-value of 0 as above.

For all candidate Bt-resistance loci that we mapped, we performed a permutation test to determine the probability that the locus overlapped with outlier sweep CLR values by chance. To do so, we randomised the position of a locus of the same size in order to generate a null distribution of overlapping mean and maximum CLR values (Supplementary Table S4).

### Calling *Kinesin-12* genotypes

To directly compare *kinesin*-12 genotypes to those reported by Benowitz *et al*. (2022), we mapped reads to the full *kinesin*-12 sequence (including untranslated regions and introns) of the LAB-S (laboratory susceptible) *H. zea* strain. This was done to rule out the potential effect of alignment errors on genotype calls and enable an unbiased comparison of genotypes with our wild-caught individuals. This is because the *H. armigera kinesin*-12 sequence differs structurally from that of *H. zea* in non-coding regions, though its position in the genome is concordant. For all 237 2019 samples, the same pipeline applied to call set 1 was used to process fastq files, map reads and call variants with HaplotypeCaller. Three different parameters were used: the LAB-S *kinesin*-12 sequence was used as the reference sequence for mapping, no more than two alternate alleles were permitted per site, and monomorphic sites were retained. When calling variants, a cut-off of 95% confidence was used (Reference genotype quality or QUAL > 13.0103 for monomorphic and polymorphic sites, respectively). Putative indels were not considered. The resulting sequences were translated using EMBOSS Transeq and aligned (along with the coding sequence of the *Bt*-resistant strain) using Mview (Madeira et al. 2022)This alignment confirmed that the *Bt*-resistant line carried at C>T mutation resulting in a premature stop codon, and allowed us to directly compare our samples.

We characterised nonsynonymous mutations of interest that could be confidently called in our samples. We reasoned that amino acid changes with novel biochemical properties are more likely to impact enzymatic function, that singleton mutations are more likely to be erroneous genotype calls, and that singleton mutations are less likely to occur at sufficient frequency to cause the putative signature of selection in the region. Therefore, we only report nonsynonymous mutations (relative to the coding region of the reference LAB-S complete sequence) that (1) produce amino acids with different biochemical properties, and (2) could be confidently called in more than one individual. Only single nucleotide polymorphisms (homozygous and heterozygous) were reported (Supplementary Table S5).

### Estimating the selection coefficient, *s*

For each site in which the CLR was calculated above, SweepFinder2 was also used to calculate Durrett and Schweinsberg’s (2004) approximation of *α*, where

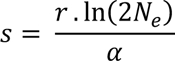

This relationship allowed us to generate an estimate of the selection coefficient *ŝ* in the sweep region on chromosome 13. To do so, we estimated the effective population size of our North American *H. zea* samples as 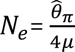 = 10.3×10^5^, using the *Drosophila melanogaster* mutation rate *μ* = 8.4×10^−8^ (Haag-Liautard et al. 2007) and 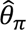 = 0.0346, calculated as the mean nucleotide diversity in 10kbp windows using pixy (Korunes & Samuk 2021). This effective population size is approximately half that of native *H. armigera* populations and consistent with previous estimates (Anderson et al. 2018). We used the mean estimate of per-base pair recombination rate for chromosome 13 (see below): *r* = 5.447×10^−8^. The mean selection coefficient within the bounds of the *CYP333B3* locus was *ŝ* = 0.0489.

### Estimating chromosome-wide recombination rates

Estimation of the selection pressure, above, required mean per-chromosome estimates of the recombination rate. We estimated recombination rates across all chromosomes using patterns of linkage disequilibrium. First, we statistically phased data across all the *H. zea* individuals we sampled for all chromosomes using Beagle v5.0 (Browning & Browning 2007, 2016). We used 50 individuals with the highest depth of coverage to estimate the population recombination parameter *ρ* across all autosomes with LDhelmet v1.9 (Chan et al. 2012). We generated likelihood lookup tables using the same theta estimator described above, with the grid of *ρ* values between 0 and 10 specified with the command -r 0.0 0.5 3.0 1.0 10.0. Following the recommendation of Chan *et al*. (26), 11 Padé coefficients were computed and, using a default block penalty of 50 and a window size of 50 SNPs, the Markov Chain Monte Carlo procedure was implemented for 10^6^ iterations with a burn-in of 10^5^ iterations. To compare observations with other data and confirm that chromosome-wide estimates of recombination rate were biologically realistic, we converted mean estimates of 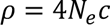 (for a recombination rate *c* per bp), to units of centimorgans per bp: 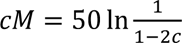. As expected, recombination rate was negatively correlated with chromosome length (Supplementary Figure S7), the gradient of this relationship was similar to that observed in *Heliconius* butterflies, and the range of recombination rates observed overlapped with those measured in both *Heliconius* and *Drosophila* (Chan et al. 2012; Martin et al. 2019).

### Modelling selection on *CYP333B3*

Taylor *et al*. (2021) genotyped *CYP333B3* from *H. zea* specimens sampled in the US at in 1998, 2002 and 2017, reporting an increase in the proportion of individuals with a derived genotype over time. We were therefore interested to know whether our estimate of *ŝ* = 0.0489 could explain these independently estimated shifts in the derived allele frequency, *q*. Starting from an observed initial allele frequency of *q*_1998_ = 0.1, we modelled the change in allele frequency in each subsequent generation as

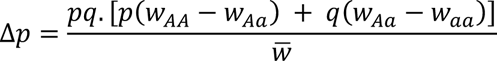

Here the reference genotypic fitness is homozygous ancestral, *i.e*.

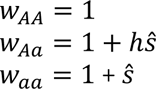

So the mean fitness is

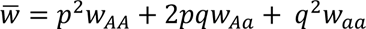

We predicted allele frequencies through time assuming a dominant-acting *CYP333B3* mutation (ℎ = 1; Figure 8B) and assuming codominance (ℎ = 0.5; Supplementary Figure S8) for a range of generation times between 2 and 10 generations per year. The biologically realistic range of generation times in this case is unlikely to be below 5 generations/yr (Hardwick 1965; Morey 2010; Parajulee et al. 2004).

Additionally, we compared our estimate *ŝ* to the selection coefficient that best explained the allele frequencies observed by Taylor *et al., ŝ_fit_*. To estimate *ŝ_fit_*, we ran 10^6^ iterations of the model with a random selection coefficient, retaining the coefficient that maximised the fit to the data (the fit was quantified as 1 minus the absolute difference between the observed and expected allele frequencies for years where allele frequencies were measured). This was done assuming complete dominance (ℎ=1), assuming codominance (ℎ=0.5), and with a random dominance coefficient to identify the combination of selection and dominance coefficients that maximised the fit to the data (Supplementary Table S6; supplementary Figure S9). The model and optimisation algorithm are available at the Github repository listed under ‘Data Availability’.

We emphasise that our aim was to determine whether our estimate of the selection coefficient was reasonable, *i.e.* whether it was in the ballpark of values that could explain independently observed allele frequency estimates. This approach is not appropriate for determining a precise estimate of the selection coefficient; four important caveats should be considered. First, we estimate a single selection coefficient over hundreds of generations. Realistically, the strength of selection imposed by pesticide exposure will vary substantially over space and time. Since the effect of selection on allele frequency depends not only on its strength but on the standing frequency, our estimate will not necessarily reflect the average strength of selection over time. Second, we compare our retrodiction with empirical data that is itself only a course estimate of the actual allele frequency through time; Taylor *et al*. sampled between 22 and 52 chromosomes, so sampling error substantially affects the observed allele frequency. Third, only three timepoints are used, and since we model Δ*p* from *q*_1998_ (to avoid any assumptions about when and how the allele first arose), we are only able to compare our estimates to two timepoints. The timepoints are, however, conveniently spaced for distinguishing between the distinct allele trajectories of dominant versus recessive mutations.

The fourth and most important caveat is that our comparison of *ŝ* and *ŝ*_1998_ is qualitative as substantially different methods are used for these estimates. Durrett and Schweinsberg’s (2004) approximation of α does not take dominance into consideration. Maynard Smith and Haigh (1974) show that dominance affects the extent to which genetic diversity is reduced at neighbouring sites through hitch-hiking, but that this effect is still highly localised. Therefore, the estimate of *s* based on SweepFinder’s α may be an over-estimation if the advantageous derived *CYP333B3* allele is completely dominant, and is not directly comparable with estimates of *s* that assume the advantageous allele is dominant. Allele frequencies modelled in Figure 8B are based on *ŝ* yet assume complete dominance of the derived allele, so they should not strictly be interpreted as expected allele frequencies. Supplementary Figure S8 shows expected allele frequencies under a consistent assumption of co-dominance and produces qualitatively similar results. Although assuming co-dominance may allow for a more consistent comparison, this assumption may be biologically unrealistic: many cases of cytochrome P450-mediated pesticide resistance involve dominance coefficients >0.5, and we expect a gain-of function mutation allowing xenobiotic metabolism to act in a dominant manner (see Discussion; Cariño et al. 1994; Han et al. 2014; Sayyed et al. 2008; Achaleke & Brévault 2010; Heckel et al. 1998). Moreover, *ŝ*_1998_ was a substantially worse fit to the empirical data when codominance was assumed – the observed rate of change early in time was far more consistent with selection on a dominant mutation (Supplementary Table S6). In this study at least, the difficult decision between directly comparable estimates and biologically realistic assumptions is resolved by the fact that *ŝ*_1998_ does not vary by more than ∼0.05 for dominance coefficients between 0.5 and 1 (see supplementary Figure S); differences between *s* estimates based on historical allele frequencies and estimates of *s* based on sweep parameters are similar irrespective of whether we assume the mutation is codominant, dominant, or anywhere in between. Therefore, despite these caveats, our estimate *ŝ* is entirely consistent with independently measured shifts in allele frequency over time.

### Testing for cryptic signatures of adaptive introgression at *CYP333B3*

We used three metrics to confirm that the sweep at *CYP333B3* was due to recent selection within *H. zea* and not the result of adaptive introgression from *H. armigera*. Adaptive introgression should result in decreased genetic differentiation from the donor species and an increased time to the most recent common ancestor (TMRCA) between homologous alleles at sites abutting the selected locus, and a decrease at the locus itself (Setter et al. 2020). By contrast, recent selection should result in an increase in genetic differentiation compared to samples collected earlier in time and a decreased TMRCA at the locus and linked sites. Therefore, we calculated *F_ST_* between *H.* zea samples from 2002 vs 2019, as well as *F_ST_* between *H. armigera* and *H. zea* sampled in 2019. This was done in both 20kbp and 100kbp windows. Next, we used Gamma-SMC (Schweiger & Durbin 2023) to estimate the TMRCA between homologous alleles at each polymorphic site on chromosome 13 within individuals sampled in 2019, using the average chromosome 13 scaled recombination rate estimate 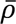 = 0.217 as a prior. This was only done within individuals (as opposed to between alleles of different individuals) so that bias resulting from potential phasing errors could be ruled out. All such analyses were carried out using call set 1.

## Data availability statement

All raw .fastq files generated here are available on the Short Read Archive: biosample accessions SAMN27502736-SAMN27502972, along with call sets as .vcf files. All custom scripts, processed data and scripts required to reproduce figures, and supplementary tables, are available at https://github.com/hlnorth/north_american_helicoverpa_zea

## Funding statement

This work was made possible through funding from a bilateral BBSRC-FAPESP grant awarded to CJ (project reference: BB/V001329/1). This work was funded in part by a USDA-APHIS Cooperative Agreement (award no. AP18PPQS&T00C204) to GS. HLN was supported by funding from the Department of Zoology, University of Cambridge.

## Conflict of interest disclosure

The authors declare no conflict of interest.

## Author contributions

HLN designed and implemented all analyses in all call sets, apart from variant calling in call set 2, under the supervision of CJ, with feedback and advice from ZF and GS. HLN wrote the manuscript text and prepared figures under the supervision of CJ with feedback from all co-authors. ZF conceived the field sampling design, collected samples, performed variant calling for call set 2, and conducted preliminary analyses not included in this publication under the supervision of GS. Other authors were integral to facilitating sample collection. We are grateful to Joana Meier and all members of the Insect Evolution and Genomics Group at Cambridge for feedback and advice on the analyses. Kyle Benowitz kindly provided the susceptible and resistant laboratory strain sequences of *Kinesin-12*, which allowed us to verify genotype calls.

## Notes

https://github.com/hlnorth/north_american_helicoverpa_zea

### Competing Interest Statement

The authors have declared no competing interest.

