## Supplementary for "Rapid evolution of pesticide resistance via adaptation and interspecific introgression in a major North American crop pest"

### Supplementary Materials

#### Supplementary Figures

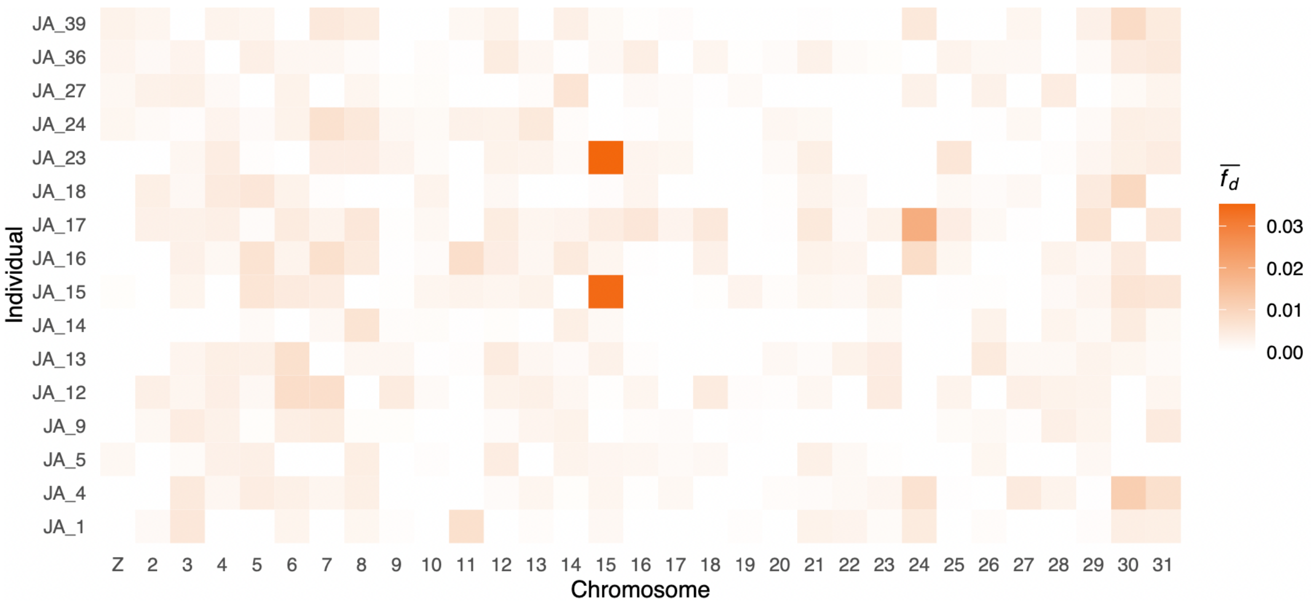


**Supplementary Figure S1: Allele sharing on chromosome 15 in two individuals.** Distribution of $\widehat{f_d}$ calculated in 20kbp windows where P1: *H. zea* sampled in 2002, P3: *H. armigera*, outgroup: *H. punctigera*. The statistic was calculated 16 times for each individual sampled in Jackson County, TX in 2019.

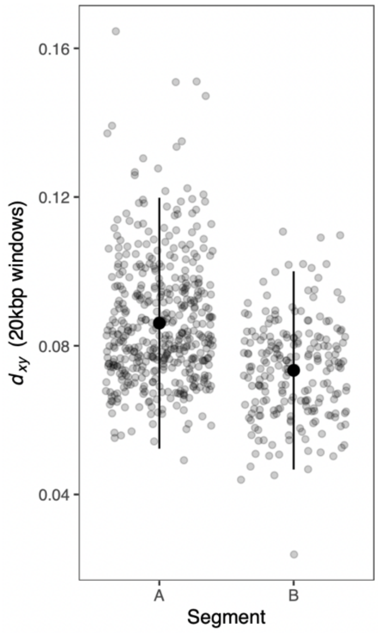


**Supplementary Figure S2: Lower genetic divergence on segment B .** Genetic divergence ($d_{xy}$), calculated in 20kbp windows, between the admixed individuals and *H. armigera* in the chromosomal segments labelled in Figure 3B.

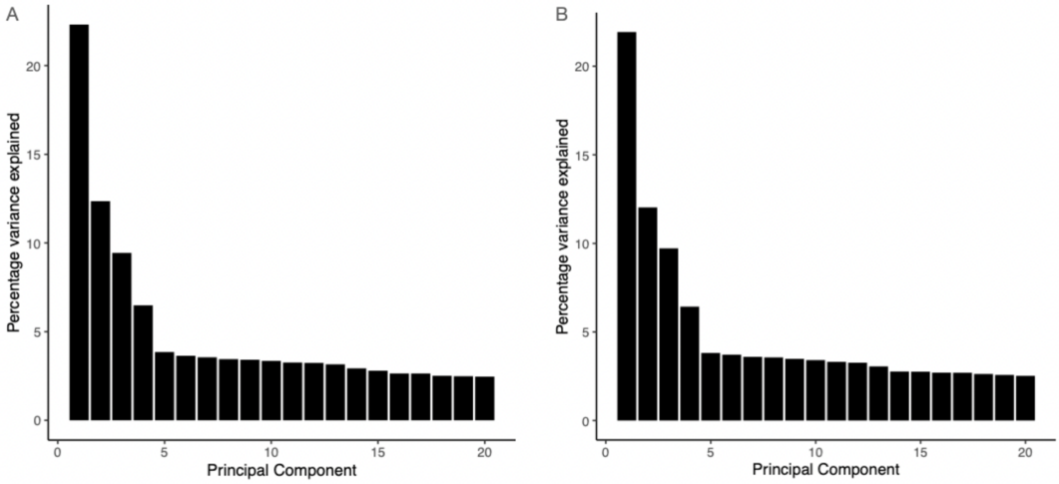


**Supplementary Figure S3: Proportion of variance explained by each principal component.** Data presented separately for chromosomal segments A and B, respectively, as show in Figure 4.

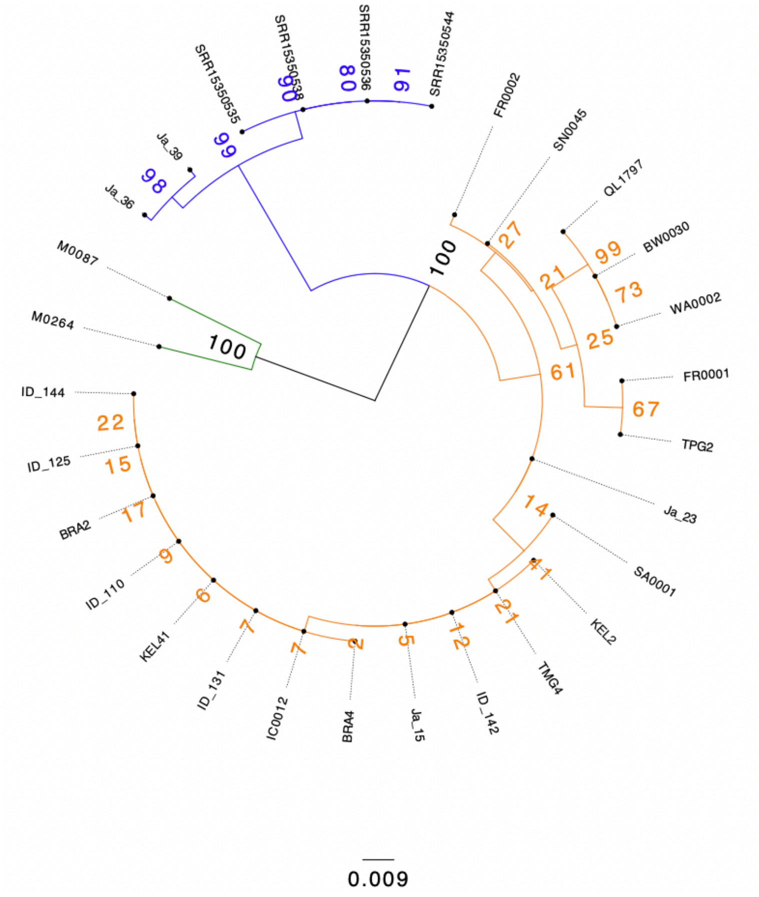


**Supplementary Figure S4: Phylogram of *CYP333B7*.** Tip labels indicate sample ID (described in Supplementary Table S2) and numbers indicate bootstrap values. This phylogram corresponds to cladogram shown in Figure 5.

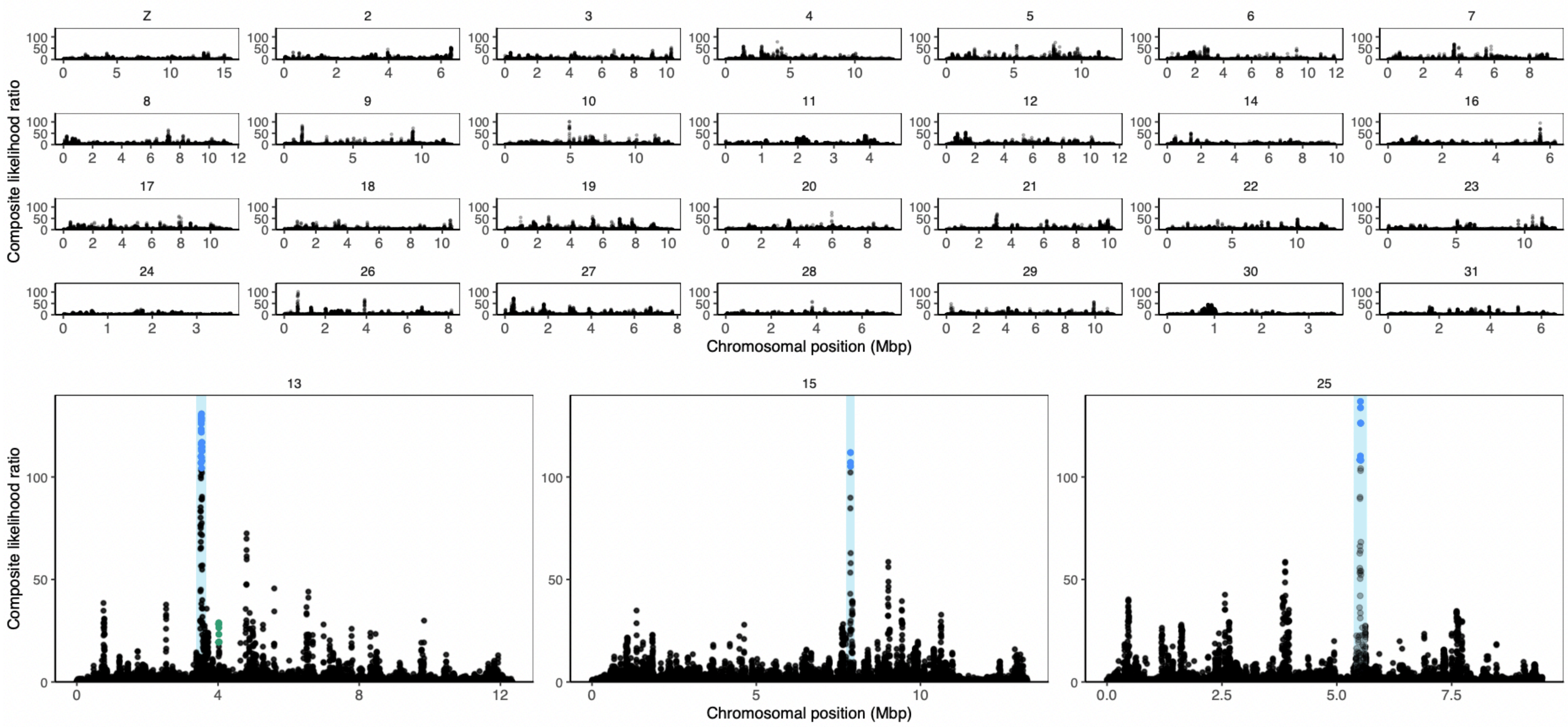


**Supplementary Figure S5: Selective Sweep composite likelihood ratio (CLR) for each chromosome.** Three chromosomes, on which selective sweeps were identified, are enlarged and highlighted. Sites in blue are in the upper 0.01st percentile of CLR values. Regions highlighted in blue shown in Figures 7 and 8. Points in green are in the upper 1st percentile and occur within the candidate Bt gene *PIK3C2A/kinesin-12*-like shown in Figure 9.

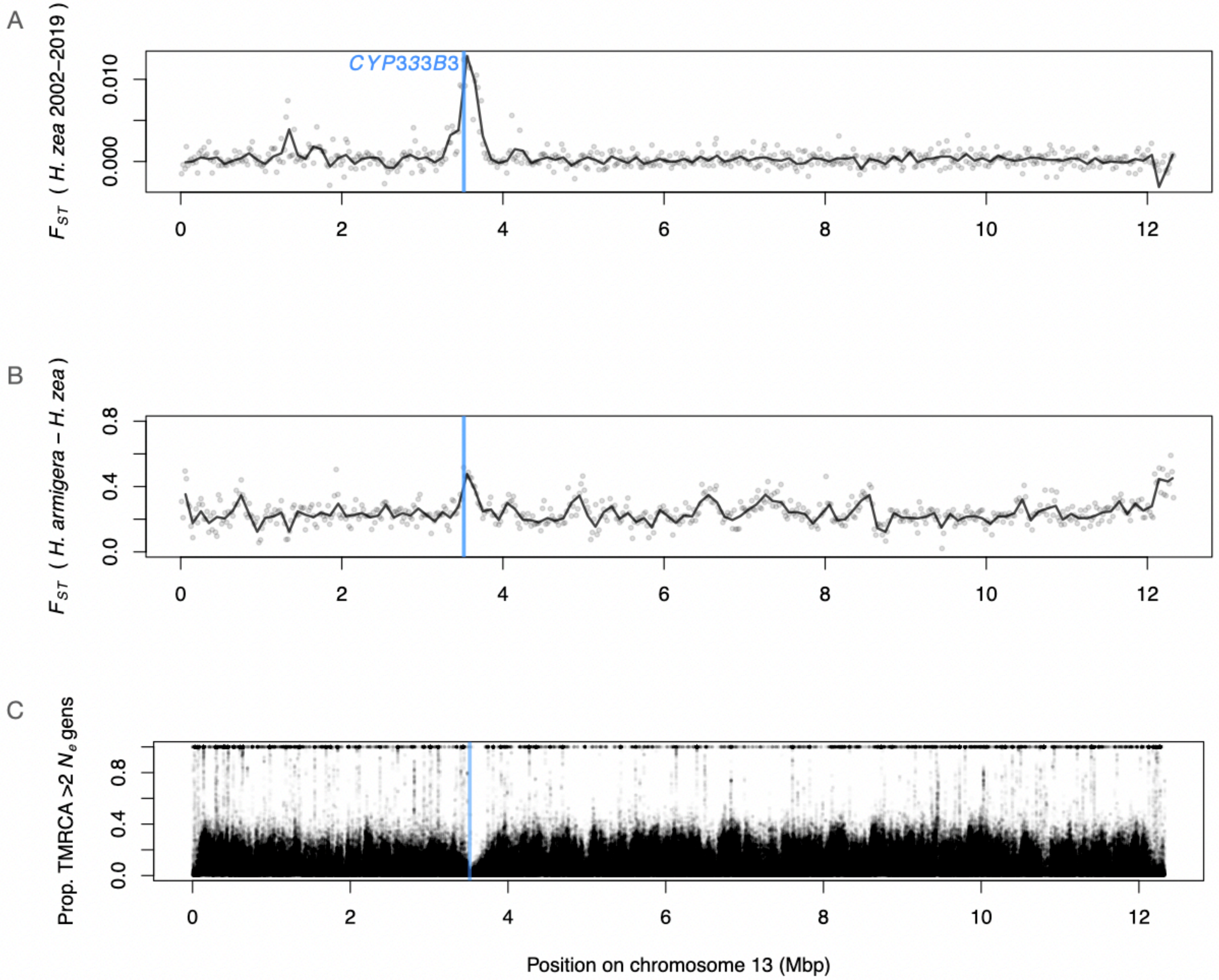


**Supplementary Figure S6: Evidence for a recent selective sweep, and against introgression, on chromosome 13. A:** Genetic differentiation ($F_{ST}$) between *H.* zea samples collected in 2002 vs. those collected in 2019, calculated in 20kbp windows (points) and 100kbp windows (lines) along chromosome 13. Blue line indicates the position of *CYP333B3*. **B:** $F_{ST}$ between *H.* zea samples collected in 2019 vs. *H. armigera*. **C:** For each polymorphic site, the proportion of *H. zea* samples collected in 2019 with an estimated time to the most recent common ancestor (TMRCA) greater than $2N_e$generations in the past between homologous alleles. Samples from 2002 were collected by Taylor *et al.* (2021).

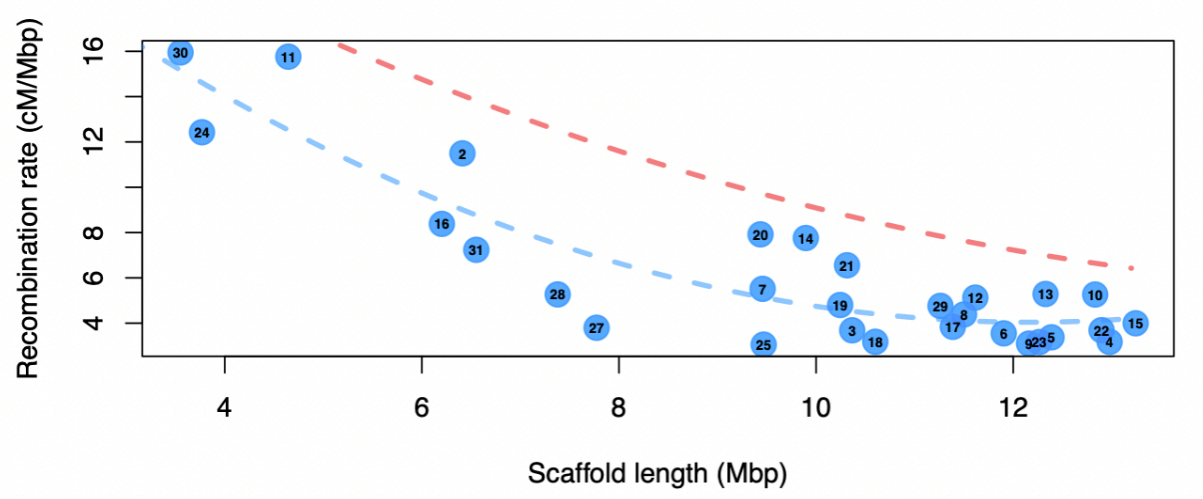


**Supplementary Figure S7: Greater estimated mean recombination rate on shorter chromosomes.** Per-chromosome recombination rate estimates against scaffold size. Blue dashed line shows model fit to data presented here. Red dashed line shows model fit to data from *Heliconius* butterflies generated by Martin *et al.* (2019).

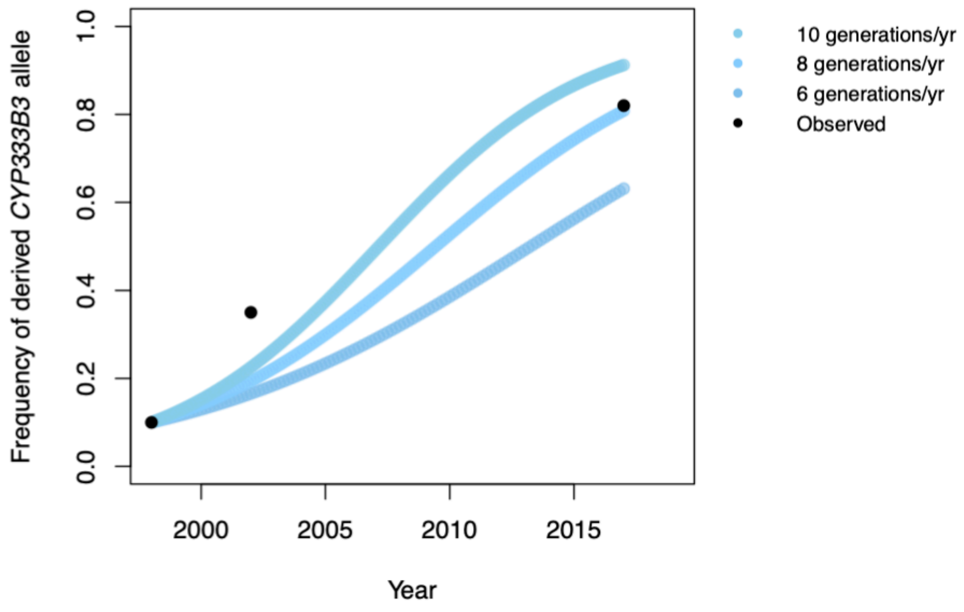


**Supplementary Figure S8: A model assuming codominance underestimates derived *CYP337B3* frequency early in the sweep.** Predicted frequency of a codominant derived *CYP333B3* allele for each generation given the estimated selection coefficient at that locus $\hat{s} = 0.0489$ assuming 6, 8 and 10 generations per year. Point colour corresponds to assumed generation time. Black points are independently estimated allele frequencies from Taylor *et al*. (2021).

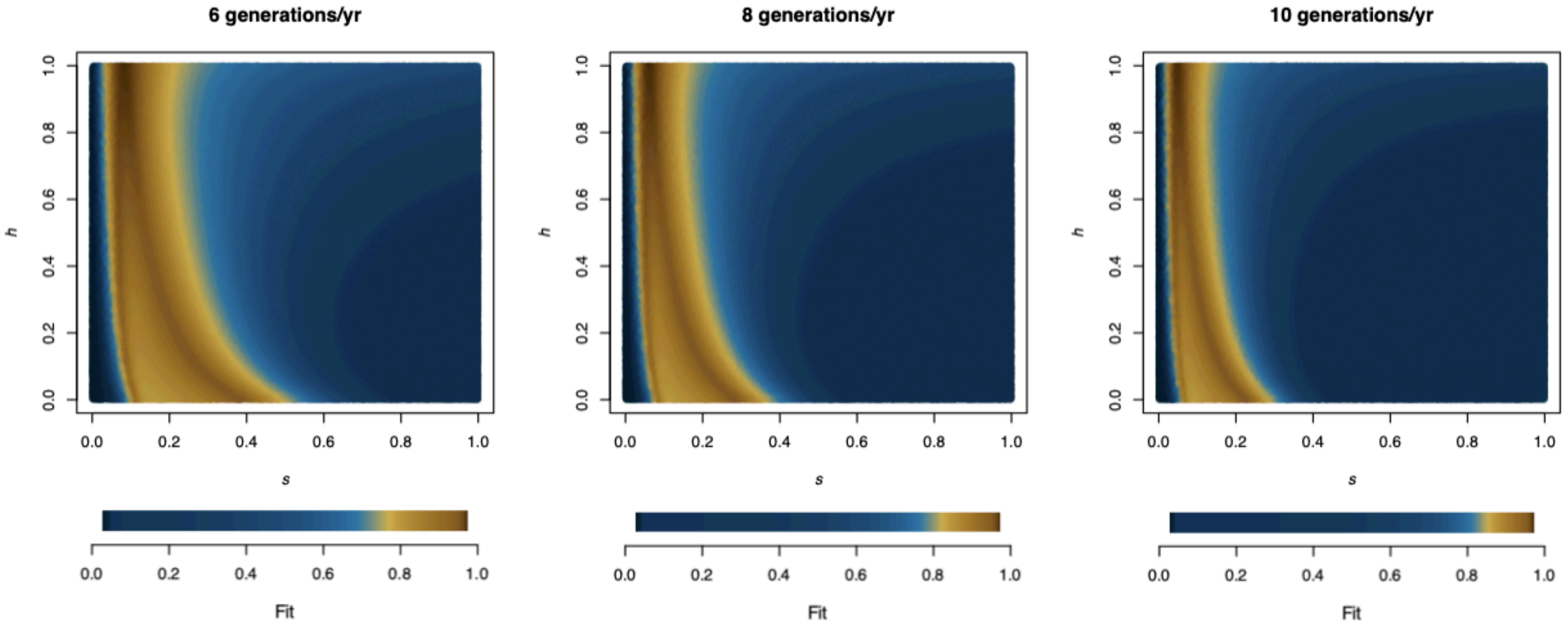


**Supplementary Figure S9: Estimates of the selection coefficient, dominance coefficient and number of generations that best explain observed allele frequencies of *CYP333B3*.** Our model of selection was run $10^6$ iterations of the model with a random selection coefficient ($s$), dominance coefficient ($h$) and one of three possible generation times. For each, the model fit was calculated. Model fit was quantified as 1 minus the absolute difference between the observed and expected allele frequencies for years where allele frequencies were measured. The parameter values that maximised fit are reported in Supplementary Table S6.

### Supplementary Tables

All supplementary tables are also available as text files, and at https://github.com/hlnorth/north_american_helicoverpa_zea

**Supplementary Figure S1: Sample sites**

| ID | location_raw | lat | lon | sample_count | source | year |
|---|---|---|---|---|---|---|
| TX-Ja | Jackson, Co., Texas (2019) | 28.94 | -96.58 | 16 | K. Crumley | 2019 |
| TX-Ma | Matagorda Co., Texas (2019) | 28.78 | -96 | 26 | K. Crumley | 2019 |
| TX-Na | Navasota, Texas (2019) | 30.383333 | -96.083333 | 24 | D. Kerns | 2019 |
| TX-Th | Thrall, Texas (2019) | 30.588611 | -97.298611 | 22 | D. Kerns | 2019 |
| TX-Wh | Wharton, Texas (2019) | 29.311667 | -96.102778 | 21 | K. Crumley | 2019 |
| AR-La | Lafayette Co., Arkansas (2019) | 33.264167 | -93.592778 | 24 | G. Lorenz | 2019 |
| AR-Ti | Tillar, Arkansas (2019) | 33.711389 | -91.453056 | 18 | G. Lorenz | 2019 |
| LA-Wb | Winnsboro, Louisiana (2019) | 32.163333 | -91.723333 | 20 | S. Brown | 2019 |
| MO-Mi | Mississippi Co., Missouri (2019) | 36.83 | -89.29 | 35 | X. Shirley | 2019 |
| NC-LM | Lees Mill Township, NC (2019) | 35.8205262 | -76.633798 | 31 | D. Reisig | 2019 |

Supplementary Figure S2: Sample metadata

See text file and https://github.com/hlnorth/north_american_helicoverpa_zea

Supplementary Figure S3: Gene annotations in sweep regions

| gene_name | chromosome | start | end | strand | annotation_source | annotation_ID | annotation_name |
|---|---|---|---|---|---|---|---|
| CYP333B3 | 13 | 3515066 | 3521020 | - | Scipio | HaOG200024 | CYP333B3_Ha |
| uncharacterised protein | 13 | 3526715 | 3531067 | - | EVM | HaOG204154 | LOC101736753BMORI:uncharacterized protein LOC101736753 |
| Cpq-like | 13 | 3550470 | 3552580 | - | EVM | HaOG204153 | BMORI:carboxypeptidase Q-like |
| TARDB43-like | 15 | 7845568 | 7847853 | + | EVM | HaOG201405 | BMORI:TAR DNA-binding protein 43-like |
| RNF186-like | 15 | 7854240 | 7857883 | - | EVM | HaOG201406 | BMORI:E3 ubiquitin-protein ligase RNF168-like |
| Hyd-like | 15 | 7858581 | 7878070 | + | EVM | HaOG201407 | BMORI:E3 ubiquitin-protein ligase hyd-like |
| PAQR3-like | 15 | 7879195 | 7881223 | - | EVM | HaOG201408 | BMORI:progestin and adipoQ receptor family member 3-like |
| toX2-like | 15 | 7882653 | 7884625 | + | EVM | HaOG201409 | BMORI:protein takeout-like isoform X2 |
| clock-like | 15 | 7885820 | 7887805 | - | EVM | HaOG201410 | BMORI:circadian clock-controlled protein-like |
| NAA40 | 25 | 5481491 | 5482145 | - | EVM | HaOG206021 | BMORI:N-alpha-acetyltransferase 40-like |
| AR-like | 25 | 5482716 | 5485238 | + | EVM | HaOG206020 | BMORI:aldose reductase-like |
| Mcm5-like | 25 | 5485331 | 5490182 | - | EVM | HaOG206019 | BMORI:DNA replication licensing factor Mcm5-like |
| dynein-like_1 | 25 | 5490930 | 5513636 | + | EVM | HaOG206018 | BMORI:dynein beta chain, ciliary-like |
| dynein-like_2 | 25 | 5513930 | 5522191 | + | EVM | HaOG206017 | BMORI:dynein beta chain, ciliary-like |
| inversin-A-like | 25 | 5522550 | 5530858 | - | EVM | HaOG206016 | BMORI:LOW QUALITY PROTEIN: inversin-A-like |
| UCP | 25 | 5538018 | 5540668 | - | EVM | HaOG206015 | BMORI:uncharacterized protein LOC101745936 |

Supplementary Figure S4: Candidate Bt gene coordinates and permutation test statistics

| name | chr | chr_start | chr_end | mean_CLR | mean_CLR_percentile | max_CLR | pval_meanLR | pval_maxLR | permutations |
|---|---|---|---|---|---|---|---|---|---|
| HGSNAT | 13 | 3972151 | 3990195 | 1.09265783 | 0.76662745 | 4.09238 | 0.268 | 0.294 | 1000 |
| liph-b | 13 | 4007535 | 4012428 | 2.175093 | 0.86089749 | 6.267544 | 0.156 | 0.098 | 1000 |
| PIK3C2A | 13 | 4012595 | 4041462 | 10.4048056 | 0.9744208 | 28.879544 | 0.024 | 0.017 | 1000 |
| kinesin-12 | 13 | 4017085 | 4018787 | 15.896281 | 0.9865347 | 19.368941 | 0.006 | 0.002 | 1000 |
| hz_G0000111 | 13 | 4042941 | 4043929 | 4.198217 | 0.92454249 | 4.198217 | NA | NA | NA |
| hz_G0000112 | 13 | 4046460 | 4054376 | 1.88253588 | 0.84322895 | 5.255188 | 0.181 | 0.145 | 1000 |
| UBE3A | 13 | 4055437 | 4064654 | 3.7718144 | 0.91605822 | 7.390298 | 0.086 | 0.109 | 1000 |
| PDE6D | 13 | 4065371 | 4066829 | 5.169933 | 0.93927816 | 5.657602 | 0.066 | 0.058 | 1000 |
| apn1 | 9 | 905656 | 913026 | 0.2207265 | 0.5277424 | 0.875827 | 0.683 | 0.643 | 1000 |
| apn4 | 9 | 916818 | 923372 | 0.0675095 | 0.38013618 | 0.261402 | 0.899 | 0.87 | 1000 |
| white | 10 | 10530120 | 10553467 | 2.50175829 | 0.8768614 | 6.243568 | 0.135 | 0.213 | 1000 |
| tspan1 | 10 | 10636587 | 10646992 | 1.8538855 | 0.84132987 | 7.403905 | 0.179 | 0.119 | 1000 |
| cad_86C | 12 | 3443579 | 3504322 | 0.69541972 | 0.69338661 | 9.173318 | 0.479 | 0.265 | 1000 |
| map4K4 | 15 | 6352455 | 6365131 | 0.55843623 | 0.66039583 | 2.14861 | 0.46 | 0.486 | 1000 |
| abcC2 | 15 | 6465002 | 6476633 | 4.18051636 | 0.92422165 | 13.601682 | 0.075 | 0.048 | 1000 |
| Cry1Ab_KZ118765 | 9 | 6911296 | 6919881 | 0.08620925 | 0.39921444 | 0.238912 | 0.889 | 0.913 | 1000 |
| Cry1A105_Cry2Ab2_KZ118015 | 9 | 10037688 | 10052829 | 0.28438907 | 0.56254152 | 1.111046 | 0.679 | 0.749 | 1000 |
| Cry1Ab_NW_018395566 | 9 | 6711005 | 6755334 | 1.83118051 | 0.83967709 | 8.825868 | 0.181 | 0.207 | 1000 |
| Cry1A105_Cry2Ab2_NW_018395399 | 9 | 8191879 | 8338153 | 0.5715646 | 0.66394444 | 10.793561 | 0.656 | 0.363 | 1000 |

**Supplementary Figure S5: Nonsynonymous genotypes identified in *Kinesin*-12**. Position refers to nucleotide in the full gene sequence including untranslated regions and noncoding regions.

| pos | WT_allele | alternate_allele | WT_AA | alternate_AA | samples_hom | samples_het | samples_alt_hom | samples_uncallable | samples_called | obs_et | exp_het | obs_hom_derived | exp_hom_derived | alt_allele_frew | allele_freq |
|---|---|---|---|---|---|---|---|---|---|---|---|---|---|---|---|
| 1130 | C | A | Q | K | 99 | 1 | 0 | 137 | 100 | 0.01 | 0.00995 | 0 | 0.000025 | 0.005 | 0.995 |
| 1374 | C | A | T | K | 75 | 20 | 1 | 138 | 96 | 0.20833333 | 0.20290799 | 0.01041667 | 0.01312934 | 0.11458333 | 0.88541667 |
| 1239 | C | T | S | L | 91 | 10 | 0 | 136 | 101 | 0.0990099 | 0.09410842 | 0 | 0.00245074 | 0.04950495 | 0.95049505 |
| 900 | A | G | Q | R | 92 | 5 | 0 | 140 | 97 | 0.05154639 | 0.05021788 | 0 | 0.00066426 | 0.0257732 | 0.9742268 |
| 1314 | A | C | D | A | 98 | 4 | 0 | 135 | 102 | 0.03921569 | 0.03844675 | 0 | 0.00038447 | 0.01960784 | 0.98039216 |
| 911 | A | T | S | C | 94 | 2 | 1 | 140 | 97 | 0.02061856 | 0.04038686 | 0.01030928 | 0.00042513 | 0.02061856 | 0.97938144 |
| 1186 | T | A | N | K | 95 | 3 | 0 | 139 | 98 | 0.03061225 | 0.03014369 | 0 | 0.00023428 | 0.01530612 | 0.98469388 |
| 926 | G | A | D | N | 90 | 2 | 0 | 145 | 92 | 0.02173913 | 0.02150284 | 0 | 0.00011815 | 0.01086957 | 0.98913044 |
| 881 | G | A | D | N | 90 | 2 | 0 | 145 | 92 | 0.02173913 | 0.02150284 | 0 | 0.00011815 | 0.01086957 | 0.98913044 |

**Supplementary Figure S5: Estimates of the selection coefficient from observed data match estimates inferred from the sweep assuming biologically realistic generation times.** $\hat{s}_{fit}$ was estimated (1) when it was allowed to co-vary with the dominance coefficient, (2) assuming complete dominance, and (3) assuming co-dominance. This was repeated for different generation times (2-10 generations per year). Each estimate is based on $10^6$ iterations of the model. The estimate of the selection coefficient from the selective sweep was 0.04894071. Biologically realistic generation times are 8-10 generations/yr. The error score is the absolute difference between the observed and expected allele frequencies for years where allele frequencies were measured. The 'fit' value in Supplementary Figure S9 is one minus the error score.

| Generations per year | s_fit_variable_h | h_fit | error_score_variable_h | s_fit_complete_dominance | error_score_complete_dominance | s_fit_codominance | error_score_codominance |
|---|---|---|---|---|---|---|---|
| 2 | 0.2857278 | 0.9999344 | 0.02323998 | 0.2870514 | 0.02273232 | 0.2146638 | 0.151069 |
| 3 | 0.182458 | 0.99954 | 0.02581397 | 0.1827668 | 0.02543528 | 0.1386256 | 0.1522696 |
| 4 | 0.1337418 | 0.9999447 | 0.02701145 | 0.1340026 | 0.026774 | 0.1023504 | 0.1528644 |
| 5 | 0.1058976 | 0.9971798 | 0.02866044 | 0.105763 | 0.02757429 | 0.08112028 | 0.1532273 |
| 6 | 0.08732526 | 0.998097 | 0.0289676 | 0.08735047 | 0.02810524 | 0.06718035 | 0.1534518 |
| 7 | 0.07375967 | 0.9993274 | 0.02972536 | 0.0743956 | 0.02848393 | 0.05732962 | 0.1536213 |
| 8 | 0.06446656 | 0.9981957 | 0.03010817 | 0.06478627 | 0.02876881 | 0.04999753 | 0.1537447 |
| 9 | 0.0572161 | 0.9992821 | 0.02961346 | 0.05737369 | 0.02898859 | 0.0443284 | 0.1538422 |
| 10 | 0.05143565 | 0.9982418 | 0.03004393 | 0.05148375 | 0.02916437 | 0.0398144 | 0.1539259 |

#### Citations (supplementary materials)

Martin SH, Davey JW, Salazar C, Jiggins CD. 2019. Recombination rate variation shapes barriers to introgression across butterfly genomes. PLoS Biol. 17:e2006288. doi: 10.1371/JOURNAL.PBIO.2006288.

Taylor KL, Hamby KA, DeYonke AM, Gould F, Fritz ML. 2021. Genome evolution in an agricultural pest following adoption of transgenic crops. Proc Natl Acad Sci U S A. 118:e2020853118. doi: 10.1073/PNAS.2020853118/SUPPL_FILE/PNAS.2020853118.SD06.XLSX.
